# PAM adenine methylation and flanking sequence regulate SaCas9 activity in bacteria

**DOI:** 10.1101/2025.08.13.670096

**Authors:** Dalton T. Ham, Tyler S. Browne, Claire Q. Zhang, Gary W. Foo, Gregory B. Gloor, David R. Edgell

## Abstract

Cas9 nucleases are the effectors of the class 2 type II CRISPR system in bacteria and function to restrict invading DNA. They can also be used with single guide RNAs (sgRNAs) as antimicrobials and genome engineering tools in bacteria, yet applications are hindered by an incomplete understanding of Cas9-target interactions. Here, we generate large-scale SaCas9/sgRNA *in vivo* bacterial activity datasets and train a machine learning model (crisprHAL) to predict SaCas9 activity. The highest predictive performance was found when downstream sequence flanking the canonical NNGRRN PAM motif at positions [+1] and [+2] was included in model training, correlating with high *in vivo* activity on sites that included T-rich di-nucleotides in the [+1] and [+2] flanking positions. Strikingly, model predictions and experimentally determined activity in pooled sgRNA experiments in *Escherichia coli* and *Citrobacter rodentium* showed an ∼10-fold reduced SaCas9 activity at sites with 5′-NNGGAT[C]-3^*′*^ PAM [+1] sequences. Cleavage assays using plasmid DNA isolated from *E. coli* inactivated for DNA adenine methyltransferase (DAM) and SaCas9/sgRNA combinations targeting sites with NNGGAT[C] PAM sequences confirmed that adenine methylation impacts SaCas9 cleavage. Moreover, ablation of a GATC DAM site in a PAM sequence enhanced SaCas9 *in vitro* activity, whereas creation of a DAM site reduced activity, providing a mechanistic link between adenine methylation and SaCas9 activity. Our results show that a general purpose machine learning architecture can provide biologically relevant insights into SaCas9-PAM interactions that can better inform activity predictions for bacterial applications. Avoidance of adenine methylated PAM sites by SaCas9 may be a mechanism of self versus non-self discrimination or reflect an evolutionary adaptation to counter methylation as an anti-restriction strategy by phage or plasmids.

## INTRODUCTION

The Cas9 RNA-guided DNA endonucleases from the class 2 type II clustered regularly interspaced palindromic repeat (CRISPR) systems are a large family of paralogous proteins that function as the effectors of a bacterial adaptive immune system to restrict invading DNA ^1–4^. Targeting of Cas9 to DNA sites requires a protospacer adjacent motif (PAM) sequence immediately adjacent to the protospacer ^5^, read out by the PAM interacting (PI) domain of Cas9^6–8^, and a CRISPR RNA (crRNA) that is complementary to the DNA protospacer ^9^. Because the PAM site is only found adjacent to crRNA target sites of invading DNA ^10^, Cas9/crRNA cleavage at similar sites in the bacterial genome is inhibited. A *trans*acting CRISPR RNA (tracrRNA) is required for processing of the crRNA transcript and for assembly of the Cas9 ribonucleoprotein complex ^9^. The crRNA and tracrRNA are genetically fused as a single guide RNA (sgRNA) for most Cas9 applications ^2^.

Cas9 nucleases are multi-domain proteins and differ from each other in coding size, the PAM sequence requirement, and the length of crRNA used for targeting ^4^. The best-studied Cas9 nuclease from *Streptococcus pyogenes* (SpCas9) has a 5′-NGG-3^*′*^ PAM requirement, and uses 20-nt crRNAs. In contrast, the Cas9 nuclease from *Staphylococcus aureus* (SaCas9), is smaller (1053 resides), requires a canonical 5′-NNGRRN-3^*′*^ PAM (hereafter NNGRRN), and uses 21- or 22-nt crRNAs ^6,11^. Structural and biochemical studies of Cas9-target DNA interactions have identified base-discriminatory contacts by the PI domain to PAM sequences that have enabled engineering of Cas9 variants that modify binding to the PAM sequence ^12–14^.

Cas9 nucleases have been adapted as novel antimicrobials and for genome engineering in bacteria ^15–19^. One critical aspect for Cas9 applications is the accurate computational prediction of activity at desired target sites while minimizing off-target toxicity ^20,21^. Most Cas9 activity predictors were developed for mammalian applications yet poorly predict sgRNA activity outside of the organism or cell line in which the training Cas9/sgRNA activity data was generated ^22^, and fail to accurately predict activity in bacteria ^20^. This issue is compounded by the fact that there are few large-scale bacterial Cas9/sgRNA activity datasets for predictive modeling. To overcome this issue, we previously generated bacterial SpCas9/sgRNA activity datasets and developed a SpCas9/sgRNA predictor, crisprHAL (crispr macHine trAnsfer Learning) that generalized activity predictions to bacteria other than *Escherichia coli* ^20^.

Here, we expand the bacterial Cas9 toolbox by generating large-scale activity datasets for SaCas9 in *E. coli* and *Citrobacter rodentium* and use them to train crisprHAL for SaCas9 predictions. In stark contrast to other characterized Cas9 paralogs, we find that di-nucleotides at positions +1 and +2 flanking the canonical SaCas9 5′-NNGRRN-3^*′*^ PAM site are critical for activity and significantly contribute to accurate activity predictions. Strikingly, we also found that SaCas9 poorly cleaves target sites flanked by a NNGGAT[C] sequence, where NNGGAT is the PAM site and [C] is at the +1 position. Finally, we show that adenine methylation at position 5 of the PAM site reduces SaCas9 cleavage activity by ∼10-fold. Our results demonstrate that choice of target site is critical for SaCas9 bacterial applications and emphasize the diversity of protein-DNA interactions in the Cas9 family of proteins. Avoidance of adenine-methylated PAM sites may be an additional mechanism for self versus non-self recognition by SaCas9, or reflect evolutionary selection against protospacer acquisition from invading phage or plasmids that use adenine methylation as an anti-restriction mechanism.

## METHODS

### Bacterial strains

*E. coli* Epi300 (F′*λ*^−^mcrAΔ(mrr-hsdRMS-mcrBC)*ϕ*80d*lacZ*Δ*M15*Δ*(lac)X74 recA1 endA1 araD139*Δ*(ara,leu)7697 galU galK rpsL* (Str^*R*^) *nupG trfA dhfr*) (Epicenter) was used for cloning the sgRNA pools. Screening sgRNA activity using a two-plasmid enrichment was done in NEB 5-alpha F′I^*q*^ *E. coli* (F^*′*^ *proA*^+^*B*^+^ *lacI*^*q*^ Δ*(lacZ)M15 zzf::Tn10* (Tet^*R*^) */fhuA2*Δ*(argF-lacZ)U169 phoA glnV44 ϕ80*Δ*(lacZ)M15 gyrA96 recA1 relA1 endA1 thi-1 hsdR17*) strain harbouring pTox. *E. coli* NK5830 (*recA arg*/F′) and *E. coli* NK5830 (recA arg /F^*′*^ zjc::Tn5; KanR *dam*) were gifts of Dr. David Haniford (Western University). *Citrobacter rodentium* DBS100 was used for screening of sgRNA activity against chromosomal targets.

### Construction of sgRNA pools

Oligonucleotides used in this study are provided in Table S1. A list of sgRNA target sites are provided in Table S2. To create pTox+KatG, a 2 kb fragment corresponding to the *Salmonella enterica* Typhimurium LT2 *katG* gene was amplified by PCR using primers DE6665 and DE6666 and cloned into pTox using Gibson Assembly and verified by whole plasmid sequencing (Supplementary Table Primer info). We identified 548 sites with NNGRRN PAM sequences on pTox+KatG. The 21- or 22-nt DNA sequence upstream of these sites was computationally extracted and used for library construction. The sgRNA library also contained 20 non-targeting sgRNAs (Table S2). Sequences that contain BsaI-HFV2 restriction sites that generate correct overhangs for Golden Gate Cloning were added to the ends of the sgRNA sequences for subsequent cloning. The sequence 5′-CCTGGTTCTTGGTCTCTCACG-3^*′*^ was added upstream of the sgRNA and 5′-GTTTTAGAGACCGCTGCCAGTTCATTTCTTAGGG-3^*′*^ was added downstream and ordered as single-strand fragments from Integrated DNA technologies (IDT DNA). Second strand synthesis was performed using 1 *µ*g of single stranded pool DNA and equimolar amounts of primer DE5224 in NEB buffer 2 (50 nM NaCl, 10 mM Tris-HCl, 10 mM MgCl_2_, 1 mM DTT, pH 7.9) by denaturing at 94°C for 5 minutes. Primers were annealed by decreasing temperature 0.1°C/second to 56°C and holding for 5 minutes, and followed by decreasing temperature 0.1°C/second to 37°C. To the annealed oligonucleotides, 1 *µ*L of Klenow polymerase (New England Biolabs) and 1 *µ*L of 10 mM dNTPs were added and incubated for 1 hour at 37°C, followed by a 20 minute incubation at 75°C before being held at 4°C. The resulting dsDNA fragments were purified using a Zymogen DNA Clean & Concentrator-5 kit following manufacturer specifications. Golden Gate cloning was used to clone the oPool and mPool into pEndo-SpCas9 and pEndo-TevSpCas9 by combining 6 pmol of oPool or mPool, 100 ng of backbone plasmid, 0.002 mg BSA, 2 *µ*L T4 DNA ligase buffer (50 mM Tris-HCl, 10 mM MgCl_2_, 1 mM ATP, 10 mM DTT, pH 7.5), 160 units T4 DNA ligase (New England Biolabs) and 20 units of BsaI-HF-V2 (New England Biolabs) with the following thermocycler conditions: 37°C for 5 min then 22°C for 5 min for 10 cycles, 37°C for 30 min, 80°C for 20 min, 12°C inf. The resulting pool was then transformed by heatshock into *E. coli* Epi300 and plated on LB plates (10 g/L tryptone, 5 g/L yeast extract, 10 g/L sodium chloride, 1% agar) supplemented with 25 mg/mL chloramphenicol and 0.2% w/v D-glucose.

To construct the sgRNA pool targeting *Citrobacter rodentium* DBS100 (Citro sgRNA), we sequenced our in-house strain at the London Regional Genomics Centre and screened a 236-kb contig for NNGRRN PAM sequences (Table S2). Sequence upstream (21-nts) was computationally extracted, with oligonucleotides ordered and cloned as described above.

### Pooled sgRNA two-plasmid enrichment experiment

A two-plasmid enrichment experiment was used to assay sgRNA activity as previously described ^23,24^. For liquid selections, 50 ng of the sgRNA plasmid pool was transformed into 50 *µ*L *E. coli* NEB 5-alpha *F* ′I^*q*^ competent cells harbouring pTox by heat shock. Cells were allowed to recover in 1 mL of non-selective 2xYT media (16 g/L, 10 g/L yeast extract, and 5 g/L NaCl) for 30 minutes at 37°C with shaking at 225 rpm. The recovery was then split and 500 *µ*L was added to 500 *µ*L of inducing 2xYT (0.04% (w/v) L-arabinose and 50 mg/mL chloramphenicol) or to 500 *µ*L of repressive 2xYT (0.4% (w/v) D-glucose and 50 mg/mL chloramphenicol) and incubated for 90 min at 37°C with shaking at 225 rpm. The two cultures were washed with 1 mL of inducing media (1x M9, 0.8% (w/v) tryptone, 1% v/v glycerol, 1 mM MgSO_4_, 1mM CaCl_2_, 0.2% (w/v) thiamine, 10 mg/mL tetracycline, 25 mg/mL chloramphenicol, 0.4 mM IPTG) or repressed media (1x M9, 0.8% (w/v) tryptone, 1% v/v glycerol, 1 mM MgSO_4_, 1mM CaCl_2_, 0.2% (w/v) thiamine, 10 mg/mL tetracycline, 25 mg/mL chloramphenicol, 0.2% (w/v) D-glucose) respectively before addition to 50 mL of the same media that was used in the wash in a 250 mL baffled flask. These cells were grown overnight at 37°C with shaking at 225 rpm. Plasmids were then isolated using the Monarch Plasmid Miniprep Kit (NEB). The sgRNA locus was then PCR amplified using primers (Table S1) containing Ilumina adapter sequence, 4 random nucleotides, 12-mer barcodes to specify the replicate, and plasmid-specific nucleotides at the 3^*′*^ end. The resulting amplicons were sent for 150 bp paired-end Illumina MiSeq sequencing at the London Regional Genomics Center (London, ON).

### *Citrobacter rodentium* depletion assay

In 10 independent reactions, 50 ng of the Citro sgRNA pool was electroporated into 100 *µ*L *C. rodentium* competent cells and allowed to recover in 1 mL of non-selective 2xYT media (16 g/L, 10 g/L yeast extract, and 5 g/L NaCl) for 30 minutes at 37°C with shaking at 225 rpm. The recovery was then split and 500 *µ*L was added to 500 *µ*L of inducing 2xYT (0.4%) (w/v) L-arabinose and 50 mg/mL chloramphenicol) or to 500 *µ*L of repressive 2xYT (0.4%) (w/v) D-glucose and 50 mg/mL chloramphenicol) and incubated for 90 min at 37°C with shaking at 225 rpm. The resulting cultures were then added to 50 mL of LB (10 g/L tryptone, 5 g/L yeast extract, 10 g/L sodium chloride) supplemented with 25 mg/mL chloramphenicol and 0.2% w/v D-glucose and grown overnight at 37°C with shaking at 225 rpm. Plasmids were isolated using the Monarch Plasmid Miniprep Kit (NEB) and the sgRNA locus was then PCR amplified using primers (Table S1) containing Ilumina adapter sequence, 4 random nucleotides, 12-mer barcodes, and plasmid-specific nucleotides at the 3′ end. The resulting amplicons were sent for 150 bp paired-end Illumina NextSeq High Output sequencing the London Regional Genomics Center (London, ON).

### SaCas9 purification

SaCas9 was purified from *E. coli* T7 express harbouring a pET-11 SaCas9 construct with a C-terminal 6x-histidine tag. Cells were cultured at 37°C in LB media until it reached an OD600 of 0.6-0.8, after which expression was induced with 1 mM IPTG at 16°C for 20 hours. After induction, cell culture was spun for 10 minutes at 4°C at 6000xg. Pellet was resuspended in binding buffer (0.5 M NaCl, 20 mM Tris – pH 8, 10 mM imidazole, 5% glycerol; 10 ml per gram of pellet) with the addition of protease inhibitors. Cells were lysed by sonication at an amplitude of 100% for 10 rounds with on and off times of 10 seconds and 20 seconds, respectively. Lysed cells were spun for an hour at 4°C at 4000xg. Protein was filtered with a 0.22*µ*M filter and was run on a HisTrap HP His tag protein purification column. The column was washed with 10 column volumes of binding buffer, and then with 10 column volumes of a 50 mM elution buffer (0.5 M NaCl, 20 mM Tris – pH 8, 50 mM imidazole, 5% glycerol). Increasing levels of imidazole (100, 200, 300, and 500 mM) were used to elute protein and fractions were concentrated using MilliporeSigma Amicon Ultra-0.5 Centrifugal Filter Units (Fisher Scientific).

### sgRNA synthesis

Synthesis of sgRNA was performed with a one-pot *in vitro* transcription reaction using HiScribe T7 High Yield RNA Synthesis Kit (NEB). In a 20*µ*L total volume, Klenow fragment (NEB), 6.67 *µ*M of sgRNA oligo, 6.67 *µ*M of universal tracr oligo, 0.125 mM of dNTPs, 10 *µ*L of 1x RiboMAX Express T7 Buffer (half of total volume), 2 *µ*L of T7 Express Enzyme Mix, and 2 *µ*L of nuclease free water were combined into a single PCR tube and incubated at 37°C for 4 hrs. RNA concentration was determined by Nanodrop.

### SaCas9/sgRNA *in vitro* cleavage assays

SaCas9 was diluted using storage buffer (50 mM Tris-HCl (pH 8), 250 mM NaCl, 1 mM DTT, 5% glycerol) to a working concentration of 140 nM of the enzyme with 280 nM sgRNA. Proteins were pre-warmed at 37°C for 10 mins and added to a pre-warmed reaction mixture consisting of 100 mM NaCl, 50 mM Tris-HCl (pH 8.0), 10 mM MgCl2, 1 mM DTT, and 5 nM pTox DNA. Reactions were stopped using 0.5 mM EDTA, 20 mg/ml RNase A, 25 mg/ml Proteinase K, and 1xD-PBS (0.9 mM CaCl2, 2.7 mM KCl, 0.5 mM MgCl2-6H20, 0.14 M NaCl, 15.2 mM Na2HPO4) and were incubated for 30 mins at 37°C. Reactions were visualized on 1% agarose gels and the observed reaction rate (*k*_*obs*_) calculated using exponential decay.

### Whole-genome and plasmid methylation analysis

Genomic DNA was extracted from *C. rodentium* DBS100 using the NEB gDNA Extraction Kit (New England Biolabs, T3010) following manufacturer’s protocol. pTox was isolated from *E. coli dam*^*+*^ or *dam*^*-*^ strains. DNA was barcoded using the Rapid Barcoding Kit V14 (Oxford Nanopore, SQK-RBK114.96) with four barcodes (RB01-RB04). Samples were loaded onto a R10.4.1. MinION flow cell (Oxford Nanopore, FLO-MIN114) and run for 24 hours. Modified bases were called and demultiplexed using Dorado (v0.9.6) with the sup,6ma,4mC 5mC model. Resulting BAM files were merged, aligned, sorted, and indexed using Samtools 1.6. Pertinent information on modified bases was extracted from the modified BAM file using Modkit (v0.4.4). Custom python scripts indexed *Citrobacter* sgRNAs and determined if GATC motifs associated with PAM sequences were called as methylated using a 75% cutoff.

### Dataset processing and curation

Illumina sequencing reads were merged using the program Usearch ^25^. These reads were then separated by their unique 12-mer barcode sequence into control and experimental condition replicates. For the *C. rodentium* depletion experiment, the barcoded replicates are from the in-vivo pre-transformation guide validation, *E. coli* Epi300 control and *C. rodentium* DBS100 experimental conditions. For the *E. coli* plasmid enrichment experiment, the barcoded replicates are from the *E. coli* Epi300 guide validation, *E. coli* NEB-5alpha with SaCas9 repressed, and *E. coli* NEB-5alpha with SaCas9 induced.

### Score calculation and model input sequence encoding

RNA sequencing data is compositional and therefore must be transformed appropriately in order to obtain meaningful statistics ^26,27^. Using ALDEx2, read counts for sgRNA features across condition replicates were transformed into posterior probabilities and then converted to linear co-ordinates using the centre-log ratio ^26^. We used the statistics relative abundance (rab.all), difference between condition (diff.btw), and difference within condition (diff.win) outputs. On-target activity scores used in model training and testing are the unmodified diff.btw scores from the enrichment data, which are expressed as log2 fold changes, and the inverse diff.btw scores from the depletion datasets.

For model training and testing, to ensure a suitable minimum dynamic range in scores, sgRNAs in the *C. rodentium* depletion dataset with control condition replicate read counts of 20 or fewer were removed. sgRNAs that were exact matches to the *E. coli* genome or had ≤3 mismatches to the *C. rodentium* genome were also removed. For the two-plasmid pTox-KatG enrichment data, all sgRNAs with direct matches to an *E. coli* genome were removed to mitigate spurious errors related to off-target, rather than on-target, activity. Model training and testing data are provided in Table S4.

Using one-hot encoding, model nucleotide sequences were converted from standard character representations ‘A’, ‘C’, ‘G’, and ‘T’ to binary matrices [1,0,0,0], [0,1,0,0], [0,0,1,0], and [0,0,0,1] respectively. This process results in 4-by-N binary matrix inputs, with N equal to the input nucleotide sequence length passed to the model.

### Model construction, training, and hyperparameter tuning

A pre-existing machine learning architecture, referred to as crisprHAL ^20^, is used as the template for model development. The crisprHAL architecture is composed of convolutional neural networks (CNN), bidirectional gated recurrent units (BGRU), and dense neural networks (DNN). These networks are arranged in a dual branch structure, sharing an initial CNN block before branching into: 1) three subsequent CNN blocks followed by 2 DNN blocks converging to a DNN of size 1, and 2) a BGRU followed by 2 DNN blocks converging to a DNN of size 1. The outputs of both branches of the model are then concatenated and passed to a final output DNN of size 1 with a linear activation function.

The 1-dimensional CNNs use 128 filters with a window size of 3 and “same” padding. CNN blocks comprise the CNN layer followed by a LeakyReLU activation function, a 1-dimensional max pooling layer with a size of 2 and “same” padding, and a dropout layer set at 0.3. The BGRU uses a size of 128, with a recurrent dropout rate of 0.2. DNN blocks comprise a DNN of size 128 or 64, for the first and second DNN layer respectively, each followed by a LeakyReLU activation function and a dropout layer set at 0.3. The previously mentioned architectural hyperparameters are unchanged from the original crisprHAL model.

The Python package “Scikit-learn” was used to split the data into training and testing sets, as well as for the generation of datasets used in 5-fold cross validation ^28^. The function “train test split” was passed a fixed state of 1 for consistent splitting of the data. All train-test splits used in the construction and testing of the model split the data into 80% training only and 20% testing only sets to avoid inadvertent contamination. Only the training data is used in 5-fold cross validation to ensure the testing data remains unseen. All model training is performed using the “Adam” optimizer, and a mean squared error loss function. Using 5-fold cross validation, we selected optimal hyperparameters for batch size, learning rate, epochs, and input sequence length.

### Performance and evaluation of models

To evaluate the performance of our models we primarily report the Spearman rank correlation coefficient and Pearson correlation coefficient. We report Spearman correlations due to its prior use in the evaluation of Cas9-sgRNA prediction models, and its lack of dependence on a linear association between data points. However, it is evident that our input data and predictions are linearly correlated. We used the Scipy stats Python package functions “spearmanr” and “pearsonr” to calculate Spearman and Pearson correlations ^29^. In-house Python scripts were used to identify mean scores for each single and di-nucleotide position relative to the PAM.

## RESULTS

### Prediction of SaCas9 activity with deep learning

To construct a SaCas9 activity prediction model, we used the existing crisprHAL architecture that leverages a dual-branch CNN (convolutional neural network) and CNN-RNN (recurrent neural network) structure ^20^ (Fig. S1). This architecture can accurately predict SpCas9 and TevSpCas9 on-target activity in different bacteria, and includes the ability to be used in an optimized transfer learning protocol. TevSpCas9 is a fusion of the I-TevI nuclease domain to SpCas9 that creates a dual cutting nuclease with an extended target site requirement as compared to SpCas9^30^; TevSaCas9 is the analogous fusion to SaCas9 (Fig. 1A) ^17^. Targeting of TevSaCas9 is dependent on the sgRNA-target site and SaCas9-PAM interactions. However, not all SaCas9/sgRNA sites are substrates for TevSaCas9 because they lack an upstream CNNNG motif in the correct spacing for cleavage by the Tev nuclease domain, but are still cleaved by SaCas9 (Fig. 1A).

**Figure 1:**
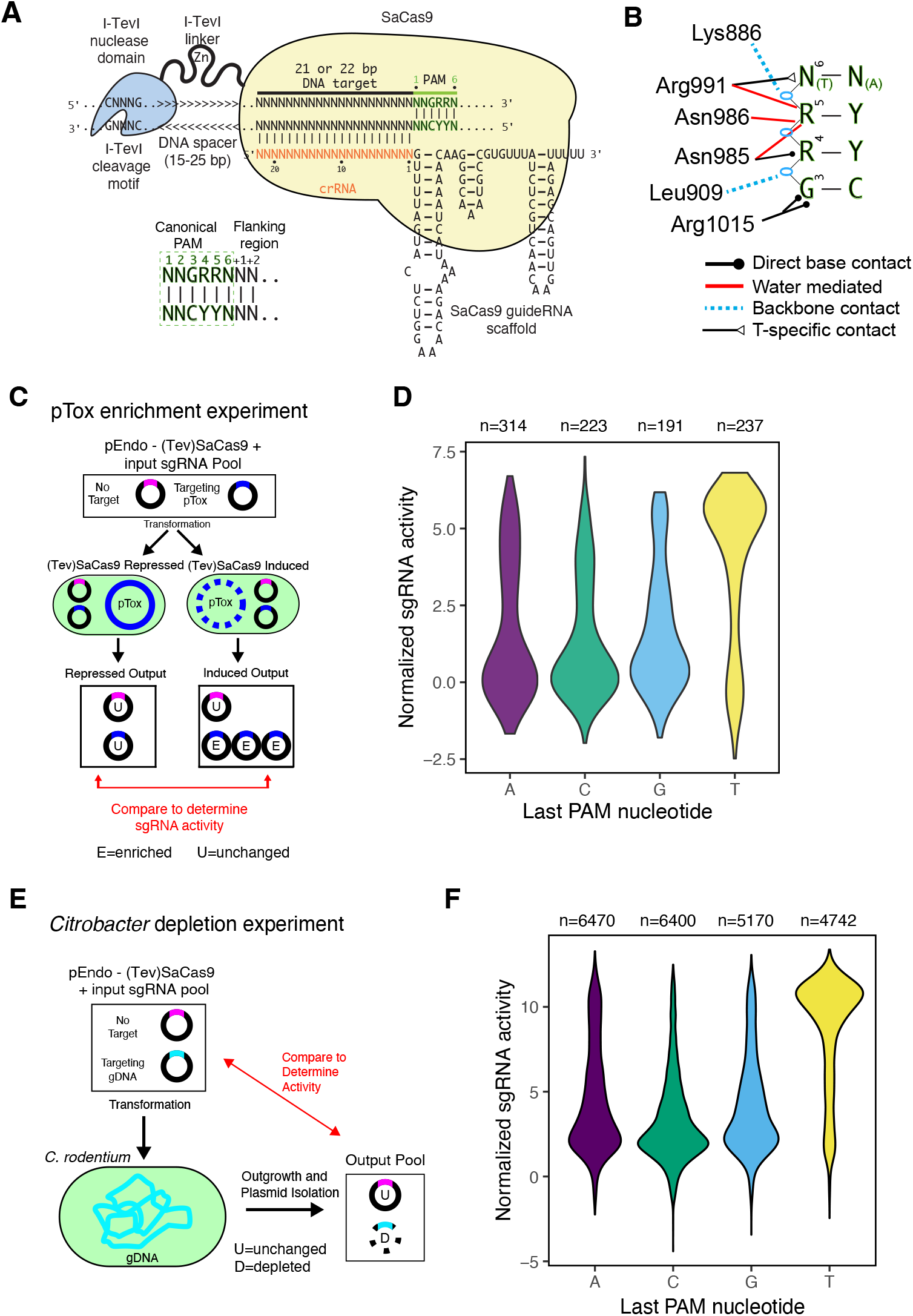
TevSaCas9/sgRNA-DNA target interactions and activity assays in bacteria. **(A)** Interactions between TevSaCas9, the sgRNA, and DNA target site (not to scale). Individual components of TevSaCas9 and the DNA target site are indicated. Zn, zinc finger in the I-TevI linker. The canonical PAM and flanking nucleotides at positions +1/+2 identified in this study as critical for SaCas9 activity are highlighted. **(B)** Summary of protein-DNA contacts in the PAM region by the PI domain of SaCas9, adapted from ref ^6^. **(C)** Schematic of the pTox enrichment experiment in *E. coli*. **(D)** Violin plot of normalized activity for sgRNAs targeted to pTox-KatG grouped by the last nucleotide of the PAM sequence. **(E)** Schematic of *C. rodentium* sgRNA depletion assay. **(F)** Violin plot of normalized activity for sgRNAs targeted to the *C. rodentium* chromosome grouped by the last nucleotide of the PAM sequence.

We used two complementary experimental methodologies to generate large-scale (Tev)SaCas9 / sgRNA activity datasets for model training. First, we used a well-established two-plasmid selection system in *E. coli* that includes a toxic plasmid expressing the CcdB DNA gyrase toxin (pTox) and a second plasmid (pEndo) that expresses (Tev)SaCas9 and an sgRNA targeting pTox (Fig. 1C) ^20,23,24^. In this assay, active (Tev)SaCas9 / sgRNA combinations will become enriched over time relative to cells expressing inactive or poorly active (Tev)SaCas9/sgRNA combinations that do not eliminate pTox and its encoded CcdB gyrase toxin. We targeted a pool of 1161 sgRNAs with 21 and 22 nts crRNA lengths to the same 548 sites on pTox (Table S2). We found no difference in activity between sgRNAs with 21- or 22-nt length crRNA regions and used 21-nt crRNAs for following experiments (Fig. S2, Table S3). Second, we used a pooled sgRNA depletion experiment that targeted 24,392 sgRNAs to the *C. rodentium* genome (Table S2). In this experiment, active (Tev)SaCas9/sgRNA combinations become depleted relative to inactive or poorly active combinations because cleavage of the bacterial chromosome cannot be repaired thereby causing DNA replication fork collapse and cell death (Fig. 1D) ^20,31^. In both datasets, we were able to identify active from inactive or poorly active (Tev)SaCas9/sgRNA combinations. We found that sgRNAs that targeted sites with PAM sequences that had a T at the last position (NNGRR**T**) showed higher activity than sites non with-T PAM sequences (Fig. 1D,E), as previously described in eukaryotic systems ^32^.

Following data curation, the *C. rodentium* depletion dataset was separated into 80% training (n = 16202) and 20% testing (n = 4051) sets. To obtain a suitable input sequence length for model training, we extended the 21-nt sgRNA target sequence upstream and downstream by 1 nt at a time, evaluating the impact of incremental nucleotide additions to model performance (Fig. 2A). Using 5-fold cross validation to assess the relative contributions of additional nucleotides, we found that the extension of the input sequence upstream of the sgRNA target (PAM distal) provided no clear improvement to model predictive performance, as measured by Spearman correlation (Fig. 2A). Conversely, inclusion of the NNGRRN PAM sequence in the model provided a large performance boost, particularly for the last 3 nucleotides (RRN) (Fig. 2A). Intriguingly, inclusion of the [+1] nucleotide downstream of the PAM provided another notable performance increase. Models constructed on each 3^*′*^ nucleotide option of the NNGRR**N** PAM supported these observations, with a notable reliance of the model on the [+1] downstream nucleotide to obtain high predictive performance (Fig. 2A). Our final TevSaCas9/sgRNA crisprHAL model obtained a high correlation for predicted and observed activities for the *C. rodentium* depletion dataset, with 0.895 Spearman (Fig. 2B) and 0.902 Pearson correlation values (Fig. S2).

**Figure 2:**
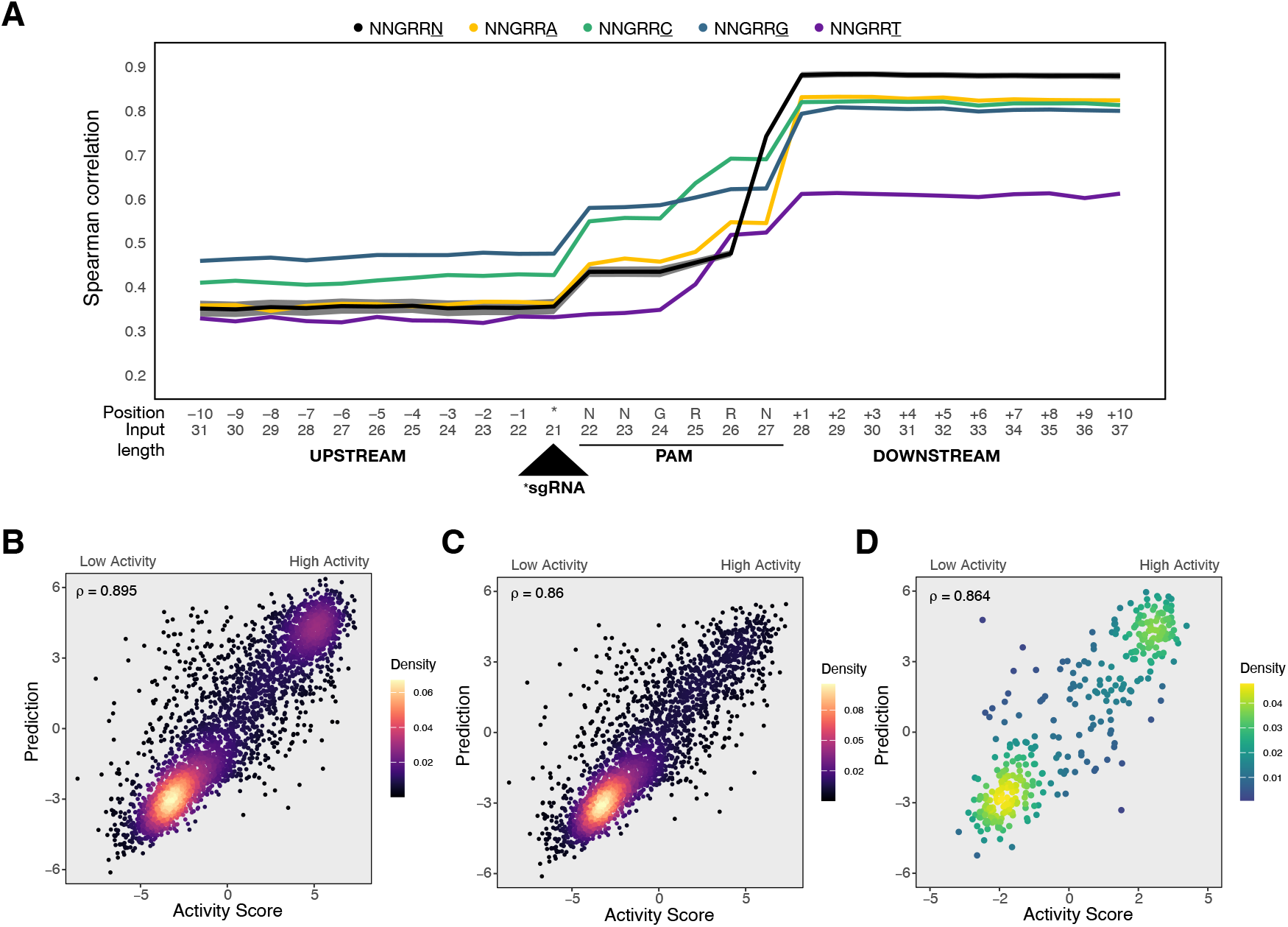
(Tev)SaCas9 target site sequence length model input optimization and model performance. **(A)** Spearman rank correlation correlation between predicted and experimentally determined on-target activity using 10 repetitions of 5-fold cross validation on the *C. rodentium* 80% training set. The primary test using all PAM sites is shown in black with the standard deviation across tests shown in dark grey. Secondary tests are separated by the final nucleotide of the NNGRRN PAM: NNGRRA (yellow), NNGRRC (green), NNGRRG (blue), and NNGRRT (purple). All tested input sequence lengths include the 21-nucleotide sgRNA target site sequence. Upstream sequence length tests extend the input sequence to include the -1 to -10 nucleotides upstream of the target site. Downstream sequence length tests extend the model input to include the NNGRRN PAM, and up to 10 nucleotides downstream of the target site. Correlations between experimentally determined and predicted on-target activity from the *C. rodentium* SaCas9 crisprHAL model on **(B)** the 20% held-out portion of the *C. rodentium* depletion dataset (n = 4051), **(C)** the NNGRRV subset of the 20% held-out portion of the *C. rodentium* depletion dataset (n = 3187), and **(D)** the *E. coli* pTox plasmid enrichment test set (n = 363).

To determine if higher activity sgRNAs targeting sites with NNGRR**T** PAMs were biasing model performance, we tested the model with a subset of sites that containing only NNGRR**V** (V=A,C,G) PAM sequences (Fig. 2C). Model performance remains high, with Spearman and Pearson correlations of 0.860 and 0.865, respectively. We also examined performance on other NNGRR**N** PAM subsets (Fig. S3). Unsurprisingly, when testing the NNGRR**T** subset, the model obtains a reduced Spearman correlation of 0.593, but a Pearson correlation of 0.802. This is likely due to a lack of diversity in the measured activity scores of sgRNAs targeting sites with NNGRR**T** PAMs relative to sites with NNGRR**V** PAM sequences. The model trained and tested on only NNGRR**V** data resulted in a test set Spearman correlation of 0.846 (Fig. S3). We note that this NNGRR**V** performance is lower than the model trained on the full NNGRR**N** dataset, indicating that model performance is not solely dependent on the identification of higher activity sgRNAs that target sites with NNGRR**T** PAMs.

To test if the crisprHAL NNGRRN model trained on the *C. rodentium* depletion dataset was transferable to other bacteria and data generated by different methods, we used the curated pTox enrichment dataset collected in *E. coli* consisting of 21-nt length sgRNAs (Table S4). We found that the NNGRRN crisprHAL *C. rodentium* model performed well on the pTox dataset, with Spearman and Pearson correlations of 0.864 and 0.891, respectively (Fig. 2D). The model also performed well on a NNGRR**V** subset of the pTox dataset (Spearman, 0.813; Pearson, 0.844) (Fig. S3).

Collectively, these data show that the crisprHAL machine learning architecture originally developed for SpCas9 can be applied to SaCas9 to generate high confidence activity predictions in different bacteria (*C. rodentium* and *E. coli*) and can be applied to datasets generated by different activity measurements (enrichment versus depletion). Surprisingly, the model identified the SaCas9 PAM motif and downstream sequence as providing significant contributions to model performance. Previous SaCas9 prediction models have focused on the contribution of the sgRNA sequence and have not considered the PAM or downstream sequence as critical for predictive modeling.

### Pyrimidine-rich di-nucleotides downstream of the PAM sequence correlate with enhanced SaCas9 activity

Inclusion of sequence downstream of the PAM site had a noticeable impact on crisprHAL model performance. To understand if the model enhancement was due to specific nucleotide preferences at downstream positions, we examined the mean activity scores for all (Tev)SaCas9/sgRNAs from the *C. rodentium* depletion dataset across 34 nts encompassing 4-nts upstream of each sgRNA binding site, the 21-nt target site and 6-nt PAM, and 8 nts downstream (Fig. 3). Plotting these scores by individual nucleotide position confirmed the striking preference for T in last position of the PAM site (NNGRR**T**), as well as a pyrimidine preference (Y=C or T) at the [+1] position downstream of the PAM (Fig. 3A). We next analyzed the contributions of di-nucleotides across the same region (Fig. 3B), revealing a strong bias towards T-rich di-nucleotides at positions N_6_[+1] and to a lesser extent at positions [+1][+2]. At these same positions, other di-nucleotides were disfavored, including GC-rich pairs. Similar preferences at individual nucleotide or di-nucleotide positions were observed with the smaller pTox enrichment dataset generated in *E. coli* (Fig. S4).

**Figure 3:**
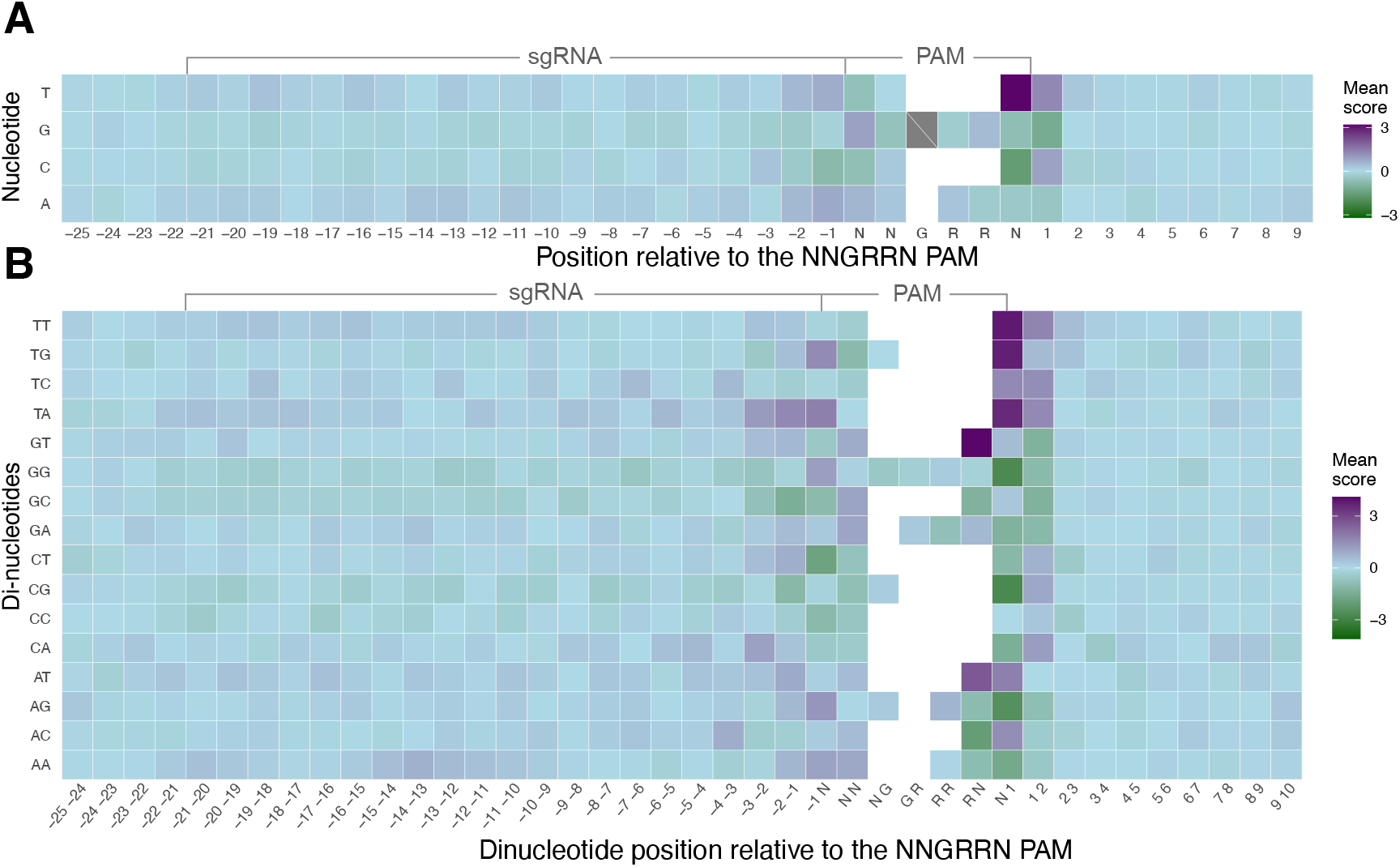
Nucleotide preference across (Tev)SaCas9 target sites. **(A)** Heatmap of mean single or **(B)** di-nucleotide activity score per position for all sgRNAs from the *Citrobacter* pooled sgRNA depletion dataset.

### Low SaCas9 activity correlates with DNA methylation

To further explore nucleotide preferences at the PAM site and flanking sequence, we focused on the last three nucleotides of the PAM (positions 4, 5, and 6) and the flanking [+1] position (NNGRRN[N]) (Fig. 4). The PI domain of SaCas9 makes base discriminatory contacts to positions 3, 4, 5 and 6 of the PAM site (Fig. 1B) ^6^, and grouping activity by these positions should reveal trends in PAM nucleotide preference. Notably, all sites with NN<u>GRR**T[N]**</u> PAM[+1] sequences had very high observed or predicted activity, except for a striking exception for sites with a NNG<u>GA**T[C]**</u> PAM[+1] (Fig. 4A,B). This analysis also showed that the majority of NNG<u>RRV**[N]**</u> PAM[+1] sequences tend to cluster at lower activity, although there are several combinations with activity equivalent to sites with a NNG<u>RRT</u> PAM, especially sites with NNG<u>RRA**[Y]**</u> PAM[+1] sequences.

**Figure 4:**
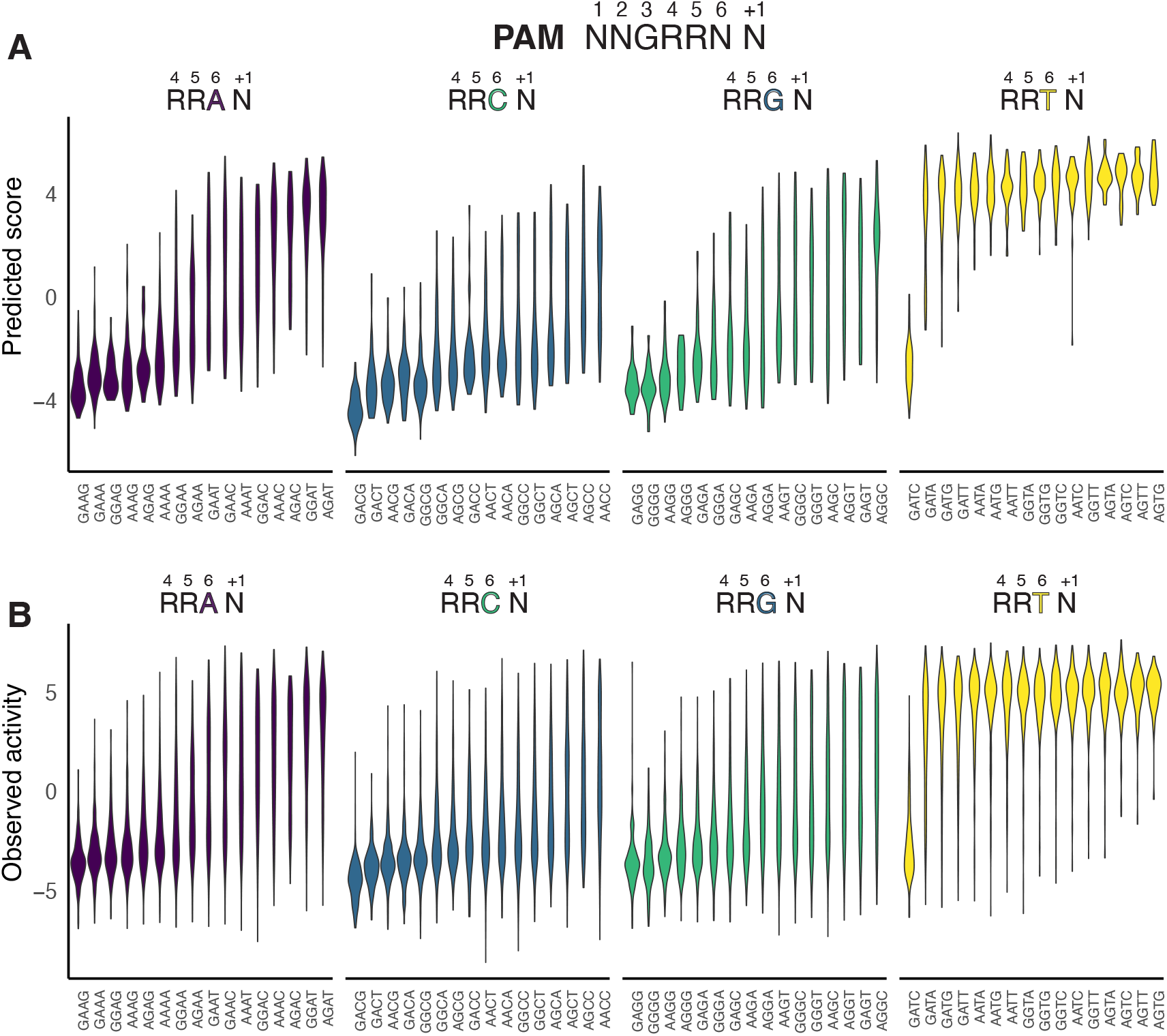
Nucleotide preference in the PAM and [+1] flanking position for sgRNA target sites in *C. rodentium*. Violin plots of **(A)** crisprHAL predicted or **(B)** observed activity for all tetranucleotide combinations at PAM positions 4,5,6, and [+1] flanking position separated by nucleotide identity at PAM position 6.

In considering why PAM[+1] sites with NNGGAT[C] sequences would be recalcitrant to SaCas9 cleavage, we noted that these sites included the sequence 5′-GATC-3^*′*^ (hereafter GATC). The sequence GATC is noteworthy because in *E. coli* and other proteobacteria it is a site for DNA adenine methyltransferase (DAM) that methylates the N6 position of A (m6A) within GATC sequences ^33^, with roles in DNA replication, mismatch DNA repair, gene regulation, and restriction-modification systems ^34,35^. PAM[+1] sites with a NNGGGT[C] sequence (where G replaces A at position 5) had high crisprHAL predicted and measured experimental activity (Fig. 4A,B). Similarly, PAM sites with NNGGAT[A], NNGGAT[T], and NNGGAT[G] sequences all had high activity, suggesting that PAM[+1] sequences containing the sequence GAT[C] with an A residue in the fifth position were responsible for the low measured and predicted SaCas9 activity. In *S. aureus* the Sau3AI restriction enzyme also recognizes GATC sites and methylation of the C (at the N5 position, m5C) within this motif protects against Sau3AI cleavage ^36,37^. Other putative and partially characterized adenine methyltransferases have been identified in *S. aureus*, and it is possible that some may be active on sequences containing a GATC motif ^38^.

We used Oxford Nanopore sequencing to map methylated DNA sequences in *C. rodentium* and to correlate methylation status with sgRNA activity (Fig 5A, B). Of the 20,253 sgRNAs tested in *C. rodentium*, 438 targeted sites with NNGGAT[C] PAM sequences and all of these were m6A methylated on both strands at GATC sites (Fig 5C, Table S5). Strikingly, sgRNA sites with NNGG*AT[C] PAM sequences had a mean observed activity of -2.70 compared to the mean activity of -0.27 for sites with NNGRRN PAM sequences. Notably, sgRNA sites with PAM sequences that differed from GATC by a single nucleotide had significantly higher activities than sites with GATC-containing PAM[+1] sequences (Fig. 5C). In particular, shifting the position of a sgRNA by a single nucleotide such that A of the GATC site was at PAM position 6 instead of position 5 increased sgRNA activity 12-fold (Fig 5B). We also considered the impact of single adenine methylation events that occurred solely in the crRNA-binding region and not in the PAM sequence; the mean observed activity at these sites was -0.32 (Fig. S5), very similar to the observed mean activity of -0.27 for sgRNAs targeting sites without any methylation in the crRNA or PAM regions. Thus, adenine methylation at sites with NNGG*AT[C] PAM sequences impacts SaCas9 activity.

**Figure 5:**
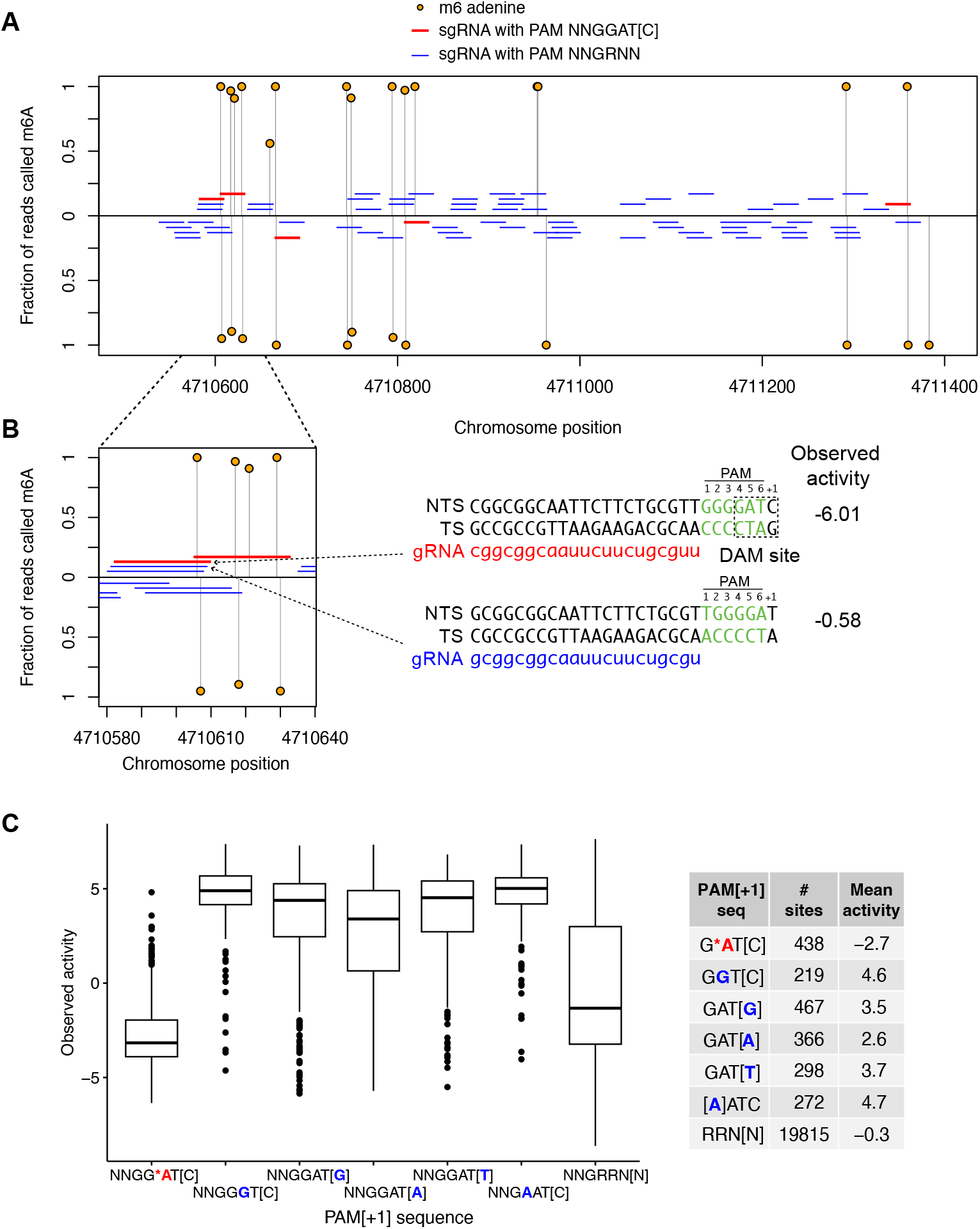
PAM adenine methylation is correlated with low sgRNA activity in *C. rodentium*. **(A)** Plot of m6A adenine methylation and sgRNA position in a representative region of the *C. rodentium* genome, with sgRNAs targeting both strands indicated. Red lines are sgRNAs with GATC sites in the PAM[+1] sequence. **(B)** Zoomed in view showing the positions of sgRNAs that target sites with and without a GATC site in the PAM[+1] sequence and their observed activity in *C. rodentium* (∼12-fold difference). Note that the two sgRNAs differ by a single nucleotide position that changes the position of the GATC site relative to the PAM sequence. **(C)** Boxplot of sgRNA activity in *C. rodentium* at target sites based on whether the sites contains a NNGG*AT[C] methylated PAM (red text), sites with single nucleotide changes in the GATC sequence (blue text), or all PAM sites. The solid horizontal line is the median point and the whiskers indicate the maximum and minimum values. Outlier data points are shown as solid circles.

We also mapped the positions of cytosine methylation relative to PAM sequences, finding that some sites with PAMs sequences that were cytosine methylated at positions 1 or 2 had low activity (Fig. S5). The bimodal distribution of activities at these sites shows that the impact on cleavage was not as consistent as for adenine methylation of PAM[+1] sequences. In support of this idea, we found that SaCas9 activity on two synthetic substrates that contained a target site with a NNGGAT[C] PAM sequence where the C at the +1 position was methylated (m5C) was equivalent to cleavage of non-methylated substrates (Fig. S6). Because SaCas9 does not make base-specific contacts to positions 1 or 2 of the PAM sequence ^6^, the reduction in cleavage is not likely to result from a disruption of specific binding due to cytosine methylation. For these reasons, we focused on characterizing the impact of adenine methylation in the PAM[+1] sequence on SaCas9 activity.

### Adenine methylation inhibits SaCas9 cleavage

To further investigate the impact of adenine methylation of PAM sequences on SaCas9 activity, we purified pTox from *E. coli dam*^+^ and *dam*^—^ strains and confirmed the DAM methylation status by DpnI digests and Oxford Nanopore sequencing. Based on the pTox pooled enrichment experiment (Fig. 1C), we picked an sgRNA with very low observed cleavage that targeted a site with a GATC PAM[+1] sequence (sgRNA_1447_) (Fig. 6A) and a high activity sgRNA that targeted a site with a GAT[G] PAM[+1] sequence (sgRNA_295_) (Fig. 6D). We found that cleavage of methylated pTox (DAM+) by purified SaCas9 and *in vitro* transcribed sgRNA_1447_ was reduced ∼3-fold relative to cleavage of non-methylated DNA (DAM-) (*k* _obs_DAM(+) 0.19 +/- 0.03 min^-1^ versus *k* _obs_DAM-0.63 +/- 0.22 min^-1^) (Fig. 6B,C). In contrast, cleavage by SaCas9/sgRNA_295_ was not affected by DAM methylation status (Fig. 6E,F).

**Figure 6:**
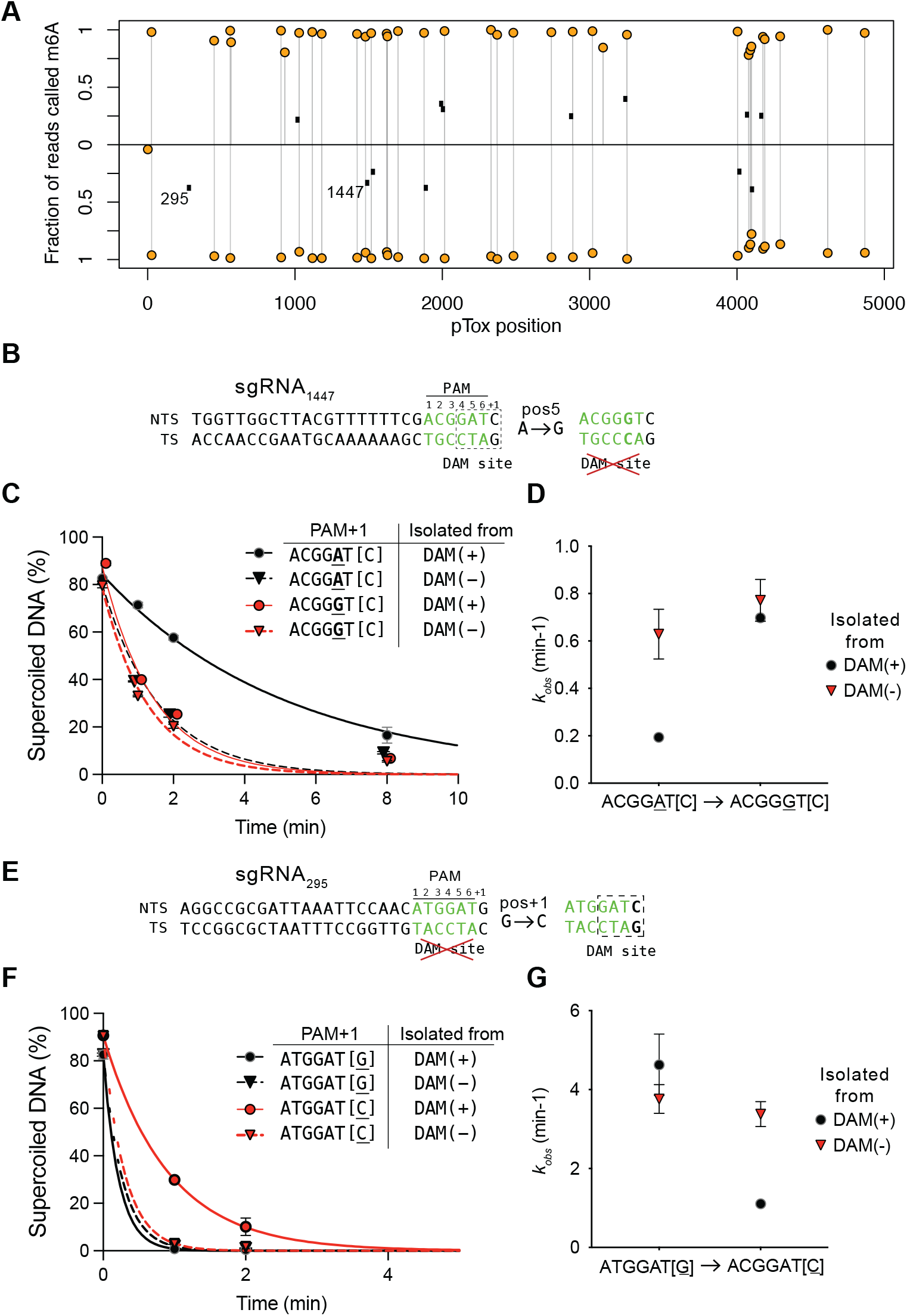
PAM adenine methylation inhibits SaCas9 activity. **(A)** Plot of methylated adenines (orange filled circles) in pTox by strand with black dots representing position of guides tested *in vitro*. sgRNA295 and 1447 that are tested below are indicated. **(B)** Sequence of pTox target site (position 1147) where SaCas9 activity is predicted to be impacted by DNA methylation (indicated by a dashed rectangle). The site-directed mutation to knockout the DAM site is indicated to the right. NTS, non-target strand; TS, target strand. **(C)** Plot of disappearance of supercoiled pTox with the wild-type or mutated target site versus time, with pTox isolated from DAM(+) or DAM(-) *E. coli* strains. **(D)** Plot of *k*_*obs*_ (min-1) calculated from plots in panel B for the different pTox plasmids isolated from DAM(+) or DAM(-) strains. **(E)** Sequence of pTox target site (position 1447) that is not methylated in the PAM+1 region, and the corresponding site-directed mutation to convert the PAM+1 to a DAM site. **(F)** and **(G)** are as for panels B and C above.

We next changed the sgRNA_1447_ PAM sequence on pTox to ACGGGT[C] from ACGGAT[C] to knockout the GATC DAM site (Fig. 6A). This change equalized the SaCas9 *in vitro* cleavage rate on pTox isolated from DAM(-) and DAM(+) strains (Fig. 6B,C). In contrast, creating a DAM site in the PAM sequence of sgRNA_295_ by changing ATGGAT[G] to ATGGAT[C] reduced SaCas9/sgRNA_295_ cleavage of methylated pTox to *k* _obs_DAM(+) 1.1 +/- 0.1 min^-1^ from *k* _obs_DAM(+) 4.6 +/- 0.8 min^-1^, a ∼4.2-fold reduction in cleavage rate. To extend these observations, we assayed 10 additional sgRNAs targeting pTox that had GATC-containing PAM[+1] sequences (Fig. 7); all showed methylation-sensitive cleavage phenotypes with mean *k*_*obs*_DAM(+) rates slower than *k*_*obs*_DAM(-) rates.

**Figure 7:**
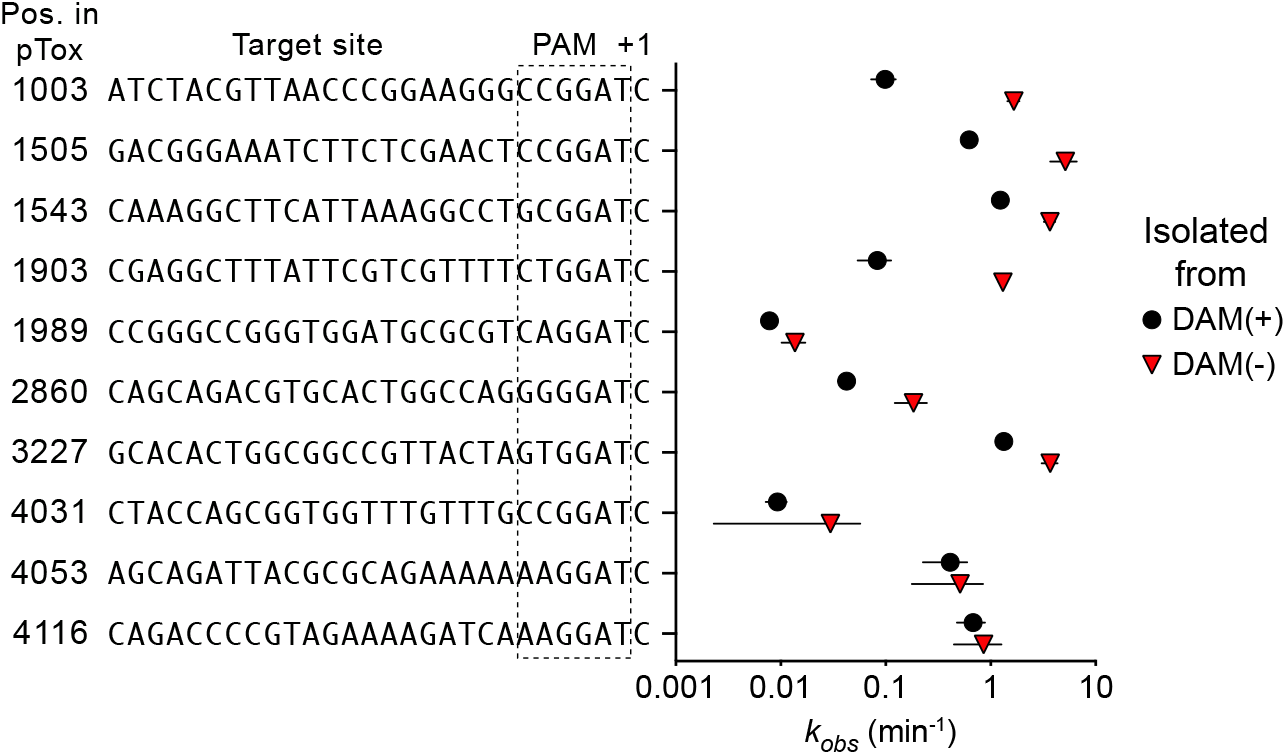
Impact of pTox methylation on SaCas9/sgRNAs *in vitro* activity. Shown is plot of *k*_*obs*_ (min^-1^) rates for 10 sgRNAs targeting different sites on pTox determined from DNA isolated from DAM(+) or DAM(-) *E. coli* strains. For all graphs points are mean of three replicates with error represented by standard deviation from the mean and plotted on log10 scale.

Collectively, this data shows a compelling agreement between machine learning predictions of SaCas9 activity, mapped epigenetic modification, and SaCas9 activity measured *in vivo* and *in vitro*. Our data indicate that adenine methylation of GATC sequences in the context of NNGGAT[C] PAM[+1] sites reduces SaCas9 activity ∼10-fold *in vivo* and ∼6-fold *in vitro* relative to unmethylated GATC sites and to sites with NNGRRN PAM sequences. Furthermore, cleavage activity *in vitro* can be manipulated by single-point mutations in PAM[+1] sequences that create or ablate GATC methylation sites, providing a mechanistic link between methylation and SaCas9 cleavage activity.

## DISCUSSION

Here, we use a combination of high-throughput biological assays, machine learning predictions of activity, methylation sequencing, and *in vitro* assays to understand factors that contribute to SaCas9 activity in bacteria. We found that the sgRNA sequence, the PAM sequence and flanking nucleotides, and adenine methylation within the PAM sequence all influence SaCas9 activity. These factors are not mutually exclusive, and do not rule out cytosine (or other) methylation as contributing to activity.

Current guidelines for SaCas9 applications suggest using a 21- or 22-nt sgRNA complementary to a DNA target site upstream of an NNGRRN PAM sequence ^11^, with some evidence that NNGRRT PAM sequences support higher activity ^32^. Available SaCas9 activity prediction models are biased towards mammalian applications and use activity data from *in vitro* methodologies or from integrated target or PAM site libraries in mammalian cells that measured activity in a fixed sequence context, rather than at sites in their native genomic context. Thus, current SaCas9 prediction tools are not trained with relevant activity data and are not suited for bacterial activity predictions. We found that using our large-scale SaCas9 activity dataset to train a general-purpose machine learning architecture ^20^, combined with appropriate data normalization and hyperparameter tuning, yielded very high prediction correlations in *E. coli* and *C. rodentium*. This finding suggests that accurate predictions could be developed for other Cas9 orthologs using the same machine learning architecture given sufficiently large, high quality activity datasets generated in bacteria.

One striking finding from our activity datasets was the influence of nucleotides downstream of the PAM sequence on SaCas9/sgRNA prediction accuracy and activity. In particular, pyrimidine-rich dinucleotides at the +1 and +2 positions downstream of the PAM, in combination with a T preference at position 6 of the PAM, had a strong positive effect on activity. The preference of SaCas9 for a T in the last position of the NNGRRN PAM sequence was noted in the initial characterization of SaCas9 PAM requirements. Structural data shows that Arg991 makes a direct contact to the O4 of thymine in position 6 of the PAM on the non-target strand ^6^, providing a rationale for a T preference. However, our observation that downstream sequence strongly impacts activity suggests that the PAM site should be revised to include NNGRRTH sequences, where H=T,C,A. One possible reason for the discrepancy between our data and previous studies could be the sequence logos methodology that was used to develop a PAM consensus sequence ^11^. Because sequence logos only takes into account the information at individual nucleotide positions it would miss the contribution of di-nucleotides at downstream positions. Interestingly, exonuclease footprinting and single-molecule studies revealed interactions of SaCas9 with flanking DNA sequence 6-bp past the PAM site ^39^. It is possible that T-rich di-nucleotides flanking the PAM site transiently stabilize the initial interaction of SaCas9 with the PAM site. We previously reported a similar observation for SpCas9^20^, where accuracy of activity predictions were increased by including downstream sequence. Interaction with DNA flanking the PAM site may be a general feature of Cas9 orthologs.

Surprisingly, we found that SaCas9 measured and predicted activity at sites with NNGGAT[C] PAM sequences was significantly lower than at target sites with other PAM sequences, including sites with NNGGGT[C] or NNGAAT[C] PAM sequences. In bacterial genomes, the sequence GATC is a methylated at the N6 position of adenine by DNA adenine methyltransferase (DAM) ^34,35^, and other methylases. In the crystal structure of SaCas9 with a DNA substrate containing a TTGAAT PAM sequence, the N6 of A5 (adenine in the 5th position) forms a water-mediated hydrogen with Asn985^6^; methylation of N6 would disrupt this hydrogen bond. In our experiments, methylation at the GATC motif within the PAM site reduces SaCas9 activity by ∼6-10 fold. We also found that PAM sites with cytosine methylation at positions 1 or 2 impacted SaCas9 activity, although this effect was not as pronounced as for adenine methylation. While the mechanistic basis of how adenine or cytosine methylation reduces activity requires further study, it is possible that adenine methylation impacts specific recognition of the A5 position by SaCas9 in the PAM sequence to destabilize or delay formation of an active R-loop required for cleavage.

Previous to our study, it was generally thought that DNA methylation had little impact on Cas9 activity ^40,41^. From an evolutionary viewpoint, it would be advantageous for Cas9 paralogs to be insensitive to DNA methylation (or other modifications), as this would be a simple strategy for invading elements to counter Cas9 activity. Interestingly, phage T4 DNA, where cytosine is replaced by glucosyl-hydroxymethylcytosine, is resistant to SpCas9 restriction if the modifications lie within the sgRNA-binding site ^42^. However, phage T4 can be restricted by SpCas9 when programmed with a sgRNA rather than with a crRNA/tracRNA combination, indicating that some high activity sgRNAs can overcome the DNA modifications. It is interesting to speculate why SaCas9 would avoid NNGGAT[C] PAM sequences - this may be an additional mechanism for self versus non-self recognition by down-regulating cleavage at methylated genomic sites that match crRNA sequences. A number of *Staphylococcal* phages are predicted to carry adenine DNA methyltransferases ^43^ and avoidance of methylated NNGGAT[C] PAM sites in phage DNA may be an evolutionary adaptation of SaCas9 to counter DNA methylation by these phages as an anti-restriction strategy. The native SaCas9 CRISPR system is not abundant in sequenced *Staphylococcal* genomes, making it difficult to correlate if naturally acquired protospacers map to phage or plasmid sequences with NNGGAT[C] PAM sites.

A recent *in vitro* profiling of PAM sequences of Cas9 paralogs identified many that could potentially contain motifs for DAM or other methyltransferases ^44^. Understanding if DNA methylation impacts activity of Cas9 orthologs would enhance on-target activity predictions for bacterial applications, particularly given that adenine and cytosine methylation are almost universal in bacteria. In contrast, it is not clear that adenine methylation impacts SaCas9 activity in mammalian systems, as evidence for adenine methylation is controversial ^45^ but may be more widespread in non-mammalian eukaryotes ^46^. We suggest that avoiding target sites with NNGGAT[C] PAM sequences would be a simple way to negate this issue.

In summary, we have shown that the combination of high-quality activity datasets and a general-purpose machine learning architecture can identify previously unrecognized DNA sequence features and epigenetic modifications that influence Cas9 activity in a biological context.

## Supporting information

DataS1

DataS3

DataS2

## Data and code availability

The Illumina and Oxford Nanopore sequencing datasets generated in this paper have been deposited in the Sequence Read Archive (BioProject ID: PRJNA1260991). The bioinformatics workflow with fixed models to replicate this study can be found at https://github.com/tbrowne5/Adenine-methylated-PAM-sequences-inhibit-SaCas9-activity. The crisprHAL prediction model and guidelines for applications can be found at https://github.com/tbrowne5/crisprHAL.

## Acknowledgments

We thank members of the Edgell and Gloor groups for reading the paper.

## Funding

This work was funded by Project Grants (PJT 159708 and PJT 191939) from the Canadian Institutes of Health Research to D.R.E. and G.B.G.

## Author contributions

Conceptualization, D.T.H., T.S.B., C.Q.Z., G.B.G., D.R.E.; methodology, D.T.H., T.S.B., G.W.F.; investigation, D.T.H., T.S.B., C.Q.Z., G.W.F.; writing – original draft, D.T.H., T.S.B. and D.R.E.; writing – review & editing, G.B.G. and D.R.E.; funding acquisition, D.R.E. and G.B.G.; supervision, D.R.E. and G.B.G.

## Declaration of interests

The authors declare no competing interests.

## Supplemental information index

Table S1. List of oligonucleotides used in this study

Table S2. List of sgRNA target sites in pTox-KatG and *C. rodentium*

Table S3. Summary of ALDEx2 outputs for sgRNA activity against pTox-KatG and *C. rodentium*

Table S4. Model training and testing datasets

Table S5. Summary of Oxford Nanopore sequencing for *C. rodentium*

Table S6. Summary table of *k*_*obs*_ rates for *in vitro* cleavage of SaCas9/sgRNA targets on pTox-KatG

Figure S1. Schematic of the crisprHAL machine learning architecture

Figure S2. Plot of sgRNA abundance versus activity for the pTox-KatG enrichmenet and *C. rodentium* depletion experiments

Figure S3. Summary plots for crisprHAL model performance on different subsets of testing data

Figure S4. Plots of nucleotide and di-nucleotide preference across (Tev)SaCas9 target sites in pTox-KatG

Figure S5. Plot of sgRNA activity for sites with single adenine or cytosine methylation in the crRNA region or PAM region in *C. rodentium*

Figure S6. SaCas9 cleavage of synthetic substrates with 5mC

## Notes

### Competing Interest Statement

The authors have declared no competing interest.

## References

1. Mojica, F. J., Díez-Villaseñor, C. s., García-Martínez, J., and Soria, E. (2005). Intervening sequences of regularly spaced prokaryotic repeats derive from foreign genetic elements. Journal of Molecular Evolution 60, 174–182.

2. Jinek, M., Chylinski, K., Fonfara, I., Hauer, M., Doudna, J. A., and Charpentier, E. (2012). A programmable dual-RNA–guided DNA endonuclease in adaptive bacterial immunity. Science 337, 816–821.

3. Koonin, E. V., and Makarova, K. S. (2019). Origins and evolution of CRISPR-Cas systems. Philosophical Transactions of the Royal Society B 374, 20180087.

4. Koonin, E. V., Makarova, K. S., and Zhang, F. (2017). Diversity, classification and evolution of CRISPR-Cas systems. Current opinion in microbiology 37, 67–78.

5. Bolotin, A., Quinquis, B., Sorokin, A., and Ehrlich, S. D. (2005). Clustered regularly interspaced short palindrome repeats (CRISPRs) have spacers of extrachromosomal origin. Microbiology 151, 2551–2561.

6. Nishimasu, H., Cong, L., Yan, W. X., Ran, F. A., Zetsche, B., Li, Y., Kurabayashi, A., Ishitani, R., Zhang, F., and Nureki, O. (2015). Crystal structure of Staphylococcus aureus Cas9. Cell 162, 1113–1126.

7. Nishimasu, H., Ran, F. A., Hsu, P. D., Konermann, S., Shehata, S. I., Dohmae, N., Ishitani, R., Zhang, F., and Nureki, O. (2014). Crystal structure of Cas9 in complex with guide RNA and target DNA. Cell 156, 935–949.

8. Jinek, M., Jiang, F., Taylor, D. W., Sternberg, S. H., Kaya, E., Ma, E., Anders, C., Hauer, M., Zhou, K., Lin, S. et al. (2014). Structures of Cas9 endonucleases reveal RNA-mediated conformational activation. Science 343, 1247997.

9. Deltcheva, E., Chylinski, K., Sharma, C. M., Gonzales, K., Chao, Y., Pirzada, Z. A., Eckert, M. R., Vogel, J., and Charpentier, E. (2011). CRISPR RNA maturation by trans-encoded small RNA and host factor RNase III. Nature 471, 602–607.

10. Gasiunas, G., Barrangou, R., Horvath, P., and Siksnys, V. (2012). Cas9–crRNA ribonucleoprotein complex mediates specific DNA cleavage for adaptive immunity in bacteria. Proceedings of the National Academy of Sciences 109, E2579–E2586.

11. Ran, F. A., Cong, L., Yan, W. X., Scott, D. A., Gootenberg, J. S., Kriz, A. J., Zetsche, B., Shalem, O., Wu, X., Makarova, K. S. et al. (2015). In vivo genome editing using Staphylococcus aureus Cas9. Nature 520, 186–191.

12. Kleinstiver, B. P., Prew, M. S., Tsai, S. Q., Nguyen, N. T., Topkar, V. V., Zheng, Z., and Joung, J. K. (2015). Broadening the targeting range of Staphylococcus aureus CRISPR-Cas9 by modifying PAM recognition. Nature Biotechnology 33, 1293–1298.

13. Walton, R. T., Christie, K. A., Whittaker, M. N., and Kleinstiver, B. P. (2020). Unconstrained genome targeting with near-PAMless engineered CRISPR-Cas9 variants. Science 368, 290–296.

14. Kleinstiver, B. P., Prew, M. S., Tsai, S. Q., Topkar, V. V., Nguyen, N. T., Zheng, Z., Gonzales, A. P., Li, Z., Peterson, R. T., Yeh, J.-R. J. et al. (2015). Engineered CRISPR-Cas9 nucleases with altered PAM specificities. Nature 523, 481–485.

15. Liu, Z., Dong, H., Cui, Y., Cong, L., and Zhang, D. (2020). Application of different types of CRISPR/Cas-based systems in bacteria. Microbial Cell Factories 19, 1–14.

16. Mayorga-Ramos, A., Zúñiga-Miranda, J., Carrera-Pacheco, S. E., Barba-Ostria, C., and Guamán, L. P. (2023). CRISPR-Cas-based antimicrobials: design, challenges, and bacterial mechanisms of resistance. ACS Infectious Diseases 9, 1283–1302.

17. Hamilton, T. A., Pellegrino, G. M., Therrien, J. A., Ham, D. T., Bartlett, P. C., Karas, B. J., Gloor, G. B., and Edgell, D. R. (2019). Efficient inter-species conjugative transfer of a CRISPR nuclease for targeted bacterial killing. Nature Communications 10, 1–9.

18. Bikard, D., Euler, C. W., Jiang, W., Nussenzweig, P. M., Goldberg, G. W., Duportet, X., Fischetti, V. A., and Marraffini, L. A. (2014). Exploiting CRISPR-Cas nucleases to produce sequence-specific antimicrobials. Nature Biotechnology 32, 1146–1150.

19. Citorik, R. J., Mimee, M., and Lu, T. K. (2014). Sequence-specific antimicrobials using efficiently delivered RNA-guided nucleases. Nature Biotechnology 32, 1141–1145.

20. Ham, D. T., Browne, T. S., Banglorewala, P. N., Wilson, T. L., Michael, R. K., Gloor, G. B., and Edgell, D. R. (2023). A generalizable Cas9/sgRNA prediction model using machine transfer learning with small high-quality datasets. Nature Communications 14, 5514.

21. Guo, J., Wang, T., Guan, C., Liu, B., Luo, C., Xie, Z., Zhang, C., and Xing, X.-H. (2018). Improved sgRNA design in bacteria via genome-wide activity profiling. Nucleic Acids Research 46, 7052–7069.

22. Konstantakos, V., Nentidis, A., Krithara, A., and Paliouras, G. (2022). CRISPR–Cas9 gRNA efficiency prediction: an overview of predictive tools and the role of deep learning. Nucleic Acids Research 50, 3616–3637.

23. Kleinstiver, B. P., Fernandes, A. D., Gloor, G. B., and Edgell, D. R. (2010). A unified genetic, computational and experimental framework identifies functionally relevant residues of the homing endonuclease I-BmoI. Nucleic Acids Research 38, 2411–2427.

24. Chen, Z., and Zhao, H. (2005). A highly sensitive selection method for directed evolution of homing endonucleases. Nucleic Acids Research 33, e154–e154.

25. Edgar, R. C. (2010). Search and clustering orders of magnitude faster than blast. Bioinformatics 26, 2460–2461. https://doi.org/10.1093/bioinformatics/btq461. doi:10.1093/bioinformatics/btq461. arXiv:http://arxiv.org/abs/ https://academic.oup.com/bioinformatics/article-pdf/26/19/2460/48857155/bioinformatics_26_19_2460.pdf

26. Fernandes, A. D., Reid, J. N., Macklaim, J. M., McMurrough, T. A., Edgell, D. R., and Gloor, G. B. (2014). Unifying the analysis of high-throughput sequencing datasets: characterizing RNA-seq, 16S rRNA gene sequencing and selective growth experiments by compositional data analysis. Microbiome 2, 1–13.

27. Gloor, G. B., and Reid, G. (2016). Compositional analysis: a valid approach to analyze microbiome high-throughput sequencing data. Canadian Journal of Microbiology 62, 692–703.

28. Pedregosa, F., Varoquaux, G., Gramfort, A., Michel, V., Thirion, B., Grisel, O., Blondel, M., Prettenhofer, P., Weiss, R., Dubourg, V., Vanderplas, J., Passos, A., Cournapeau, D., Brucher, M., Perrot, M., and Duchesnay, E. (2011). Scikit-learn: Machine learning in Python. Journal of Machine Learning Research 12, 2825–2830.

29. Virtanen, P., Gommers, R., Oliphant, T. E., Haberland, M., Reddy, T., Cournapeau, D., Burovski, E., Peterson, P., Weckesser, W., Bright, J., van der Walt, S. J., Brett, M., Wilson, J., Millman, K. J., Mayorov, N., Nelson, A. R. J., Jones, E., Kern, R., Larson, E., Carey, C. J., Polat, I., Feng, Y., Moore, E. W., VanderPlas, J., Laxalde, D., Perktold, J., Cimrman, R., Henriksen, I., Quintero, E. A., Harris, C. R., Archibald, A. M., Ribeiro, A. H., Pedregosa, F., van Mulbregt, P., and SciPy 1.0 Contributors (2020). SciPy 1.0: fundamental algorithms for scientific computing in python. Nature Methods 17, 261–272.

30. Wolfs, J. M., Hamilton, T. A., Lant, J. T., Laforet, M., Zhang, J., Salemi, L. M., Gloor, G. B., Schild-Poulter, C., and Edgell, D. R. (2016). Biasing genome-editing events toward precise length deletions with an RNA-guided TevCas9 dual nuclease. Proceedings of the National Academy of Sciences 113, 14988–14993.

31. Cui, L., and Bikard, D. (2016). Consequences of Cas9 cleavage in the chromosome of Escherichia coli. Nucleic Acids Research 44, 4243–4251.

32. Xie, H., Tang, L., He, X., Liu, X., Zhou, C., Liu, J., Ge, X., Li, J., Liu, C., Zhao, J. et al. (2018). SaCas9 requires 5-NNGRRT-3 PAM for sufficient cleavage and possesses higher cleavage activity than SpCas9 or FnCpf1 in human cells. Biotechnology Journal 13, 1700561.

33. Hattman, S., Brooks, J. E., and Masurekar, M. (1978). Sequence specificity of the P1 modification methylase (m·Eco P1) and the DNA methylase (m· eco dam) controlled by the Escherichia coli dam gene. Journal of Molecular Biology 126, 367–380.

34. Sánchez-Romero, M. A., Cota, I., and Casadesús, J. (2015). DNA methylation in bacteria: from the methyl group to the methylome. Current Opinion in Microbiology 25, 9–16.

35. Low, D. A., Weyand, N. J., and Mahan, M. J. (2001). Roles of DNA adenine methylation in regulating bacterial gene expression and virulence. Infection and Immunity 69, 7197–7204.

36. Sussenbach, J., Monfoort, C., Schiphof, R., and Stobberingh, E. (1976). A restriction endonuclease from Staphylococcus aureus. Nucleic Acids Research 3, 3193–3202.

37. Streeck, R. E. (1980). Single-strand and double-strand cleavage at half-modified and fully modified recognition sites for the restriction nucleases Sau3a and Taqi. Gene 12, 267–275.

38. Liu, J.-H., Zhang, Y., Zhou, N., He, J., Xu, J., Cai, Z., Yang, L., and Liu, Y. (2024). Bacmethy: A novel and convenient tool for investigating bacterial DNA methylation pattern and their transcriptional regulation effects. Imeta 3, e186.

39. Zhang, S., Zhang, Q., Hou, X.-M., Guo, L., Wang, F., Bi, L., Zhang, X., Li, H.-H., Wen, F., Xi, X.-G. et al. (2020). Dynamics of Staphylococcus aureus Cas9 in DNA target Association and Dissociation. EMBO reports 21, e50184.

40. Hsu, P. D., Scott, D. A., Weinstein, J. A., Ran, F. A., Konermann, S., Agarwala, V., Li, Y., Fine, E. J., Wu, X., Shalem, O. et al. (2013). DNA targeting specificity of RNA-guided Cas9 nucleases. Nature Biotechnology 31, 827–832.

41. Jiang, F., and Doudna, J. A. (2017). CRISPR–Cas9 structures and mechanisms. Annual Review of Biophysics 46, 505–529.

42. Bryson, A. L., Hwang, Y., Sherrill-Mix, S., Wu, G. D., Lewis, J. D., Black, L., Clark, T. A., and Bushman, F. D. (2015). Covalent modification of bacteriophage T4 DNA inhibits CRISPR-Cas9. MBio 6, 10–1128.

43. Ulrich, R. J., Podkowik, M., Tierce, R., Irnov, I., Putzel, G., Samhadaneh, N. M., Lacey, K. A., Boff, D., Morales, S. M., Makita, S. et al. (2025). Prophage-encoded methyltransferase drives adaptation of community-acquired methicillin-resistant Staphylococcus aureus. The Journal of Clinical Investigation. doi:10.1172/JCI177872.

44. Gasiunas, G., Young, J. K., Karvelis, T., Kazlauskas, D., Urbaitis, T., Jasnauskaite, M., Grusyte, M. M., Paulraj, S., Wang, P.-H., Hou, Z. et al. (2020). A catalogue of biochemically diverse CRISPR-Cas9 orthologs. Nature Communications 11, 5512.

45. Douvlataniotis, K., Bensberg, M., Lentini, A., Gylemo, B., and Nestor, C. E. (2020). No evidence for DNA N 6-methyladenine in mammals. Science Advances 6, eaay3335.

46. Kong, Y., Cao, L., Deikus, G., Fan, Y., Mead, E. A., Lai, W., Zhang, Y., Yong, R., Sebra, R., Wang, H. et al. (2022). Critical assessment of DNA adenine methylation in eukaryotes using quantitative deconvolution. Science 375, 515–522.

