## Supplemental Data for "PAM adenine methylation and flanking sequence regulate SaCas9 activity in bacteria"

#### Supplemental information index

- Figure S1. Schematic of the crisprHAL machine learning architecture
- Figure S2. Plot of sgRNA abundance versus activity for the pTox-KatG enrichmenet and *C. rodentium* depletion experiments
- Figure S3. Summary plots for crisprHAL model performance on different subsets of testing data
- Figure S4. Plots of nucleotide and di-nucleotide preference across (Tev)SaCas9 target sites in pTox-KatG
- Figure S5. Plot of sgRNA activity for sites with single adenine or cytosine methylation in the crRNA region or PAM region in *C. rodentium*
- Figure S6. SaCas9 cleavage of synthetic substrates with 5mC
- Table S1. List of oligonucleotides used in this study
- Table S2. List of sgRNA target sites in pTox-KatG and *C. rodentium* [1]
- Table S3. Summary of ALDEx2 outputs for sgRNA activity against pTox-KatG and *C. rodentium* [1]
- Table S4. Model training and test datasets [1]
- Table S5. Summary of Oxford Nanopore sequencing for *C. rodentium* [1]
- Table S6. Summary table of $k_{obs}$ rates for *in vitro* cleavage of SaCas9/sgRNA targets on pTox-KatG
- Data S1. GenBank file of pTox+KatG [1]
- Data S2. GenBank file of pEndo-TevSaCas9 [1]
- Data S3. GenBank file of pEndo-SaCas9 [1]
- Data S4. Fasta file of *C. rodentium* contig [1]

[1]File uploaded separately

### Supplementary Figures

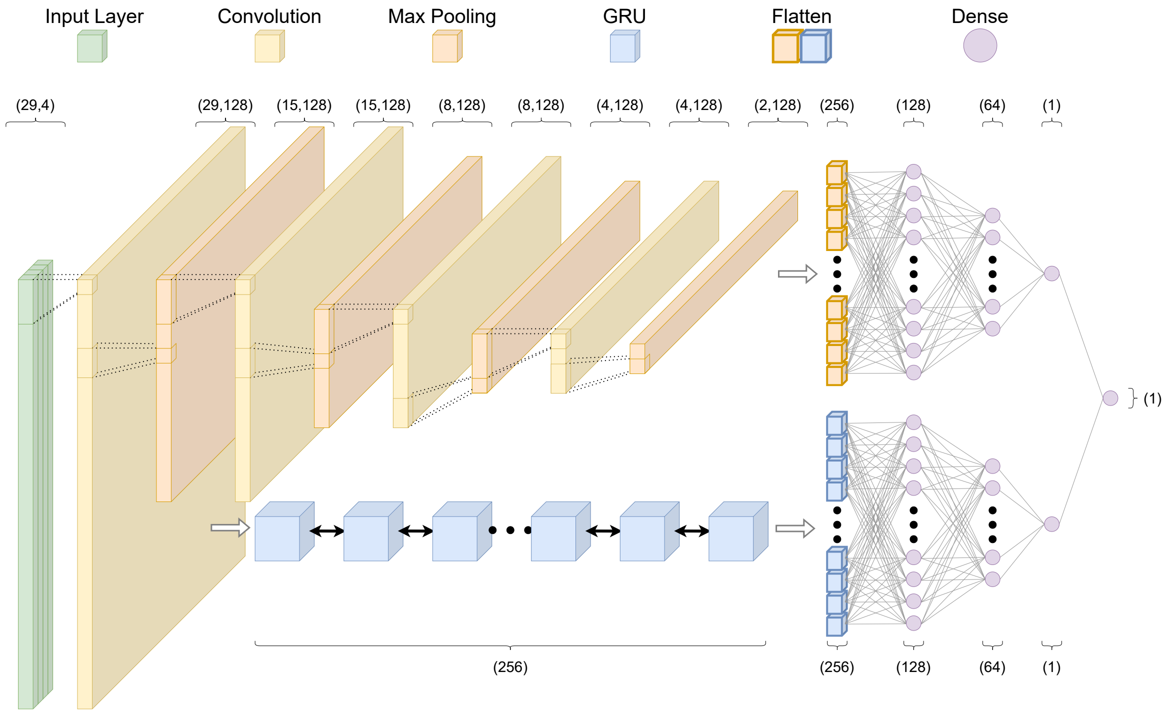


Figure S1: crisprHAL model architecture. A one-hot encoded input nucleotide sequence (input layer, green box) is passed through the dual branch CNN and bi-directional GRU RNN structure, each with subsequent dense layers, resulting in a final output prediction of on-target activity.

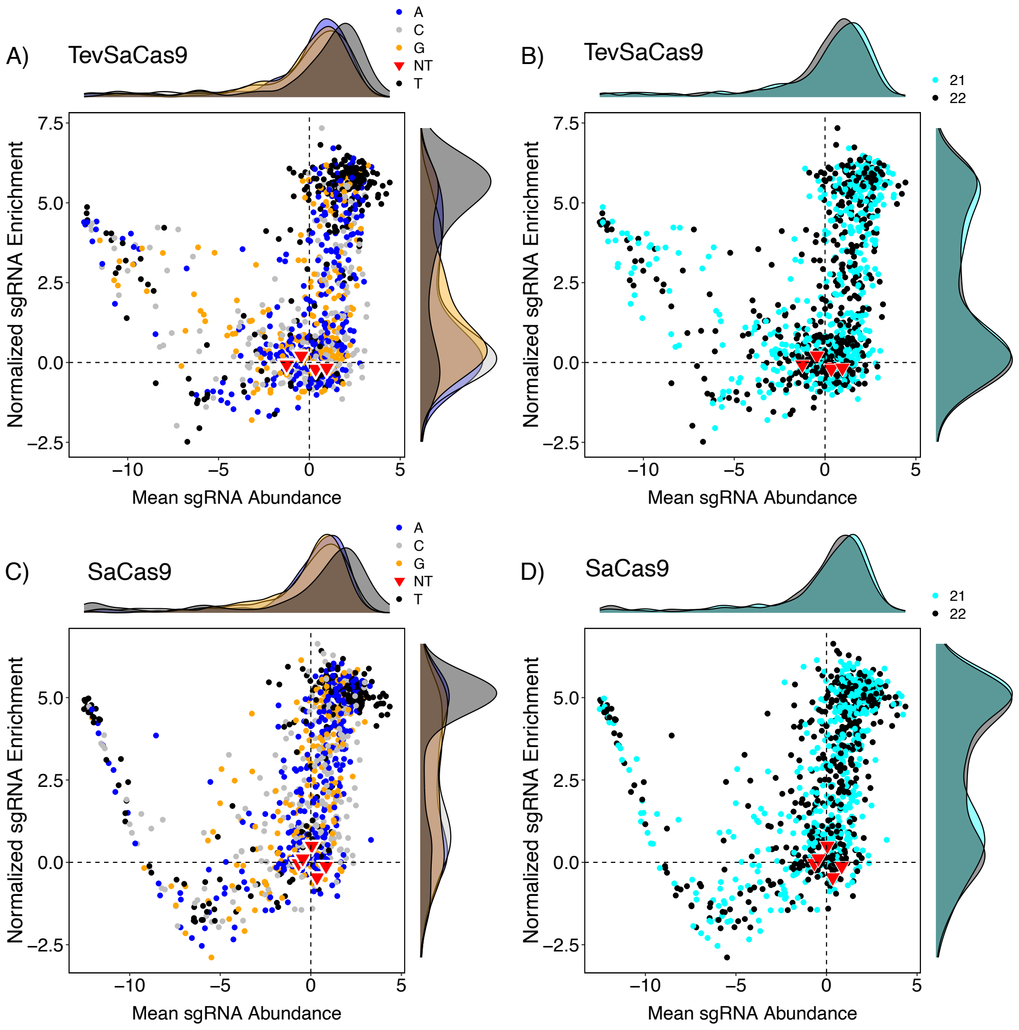


Figure S2: Enrichment experiment for the (Tev)SaCas9/sgRNA pool targeting pTox-KatG. Shown are the plots of normalized sgRNA enrichment versus mean sgRNA abundances for a pool targeting pTox+KatG with TevSaCas9 **(A,B)** and SaCas9 **(C,D)**. In panels **(A and C)** The sgRNAs were separated according to the last nucleotide in 5′-NNGRRN-3′ PAM sequence: C (grey), A (blue), T (black), G (orange) and non-targeting (red). In panels **(B and D)**, the sgRNA are separated by length: 21 nt (cyan), 22 nt (black), and non-targeting (red).

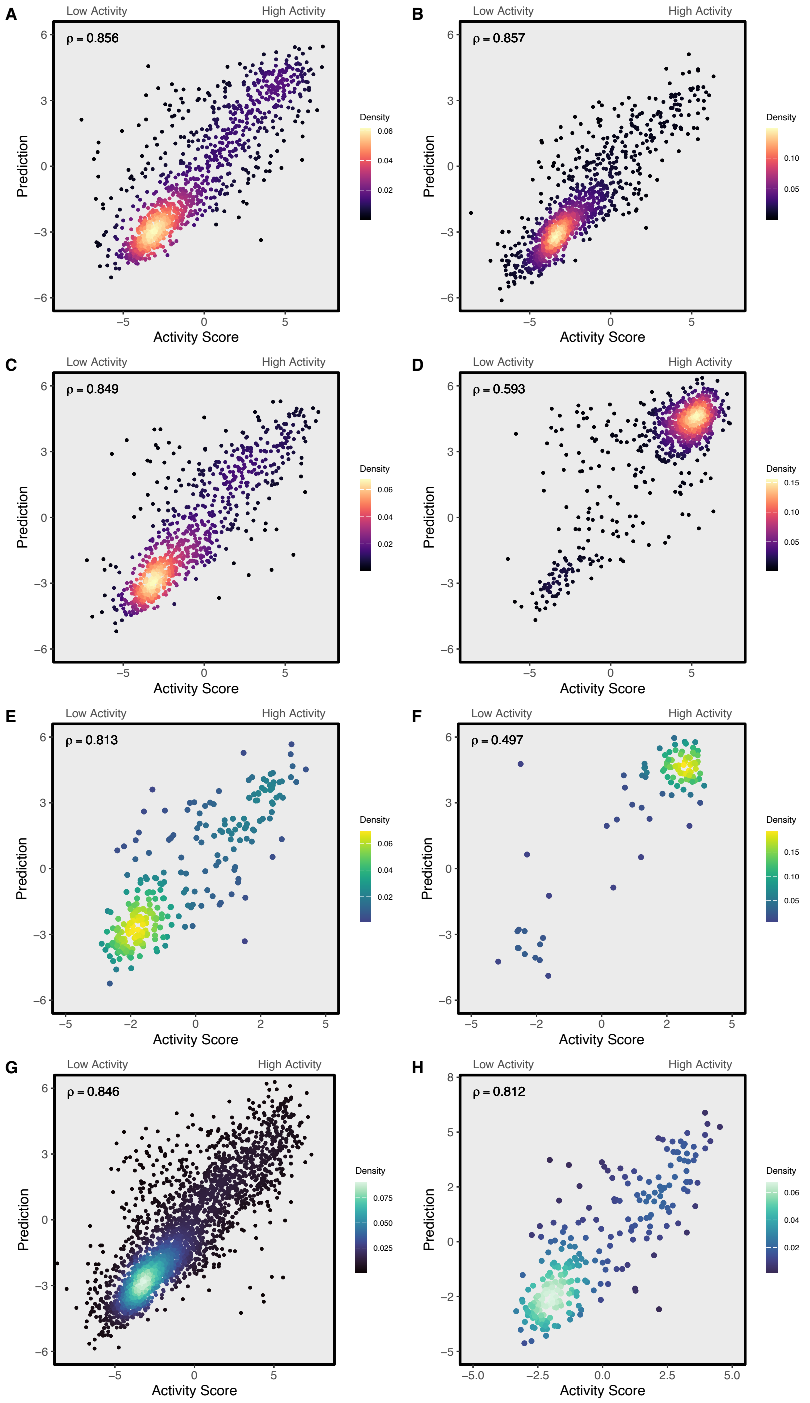


Figure S3: Predictions and Spearman correlations from the final SaCas9 crisprHAL model on subsets of the *C. rodentium* test set for sites with the following PAM sequences (A) NNGRRA, (B) NNGRRC, (C) NNGRRG, and (D) NNGRRT; and subsets of the pTox-KatG test set with the PAM sequences (E) NNGRRV and (F) NNGRRT. Predictions and Spearman correlations from the NNGRRV PAM SaCas9 crisprHAL model on NNGRRV subsets of the (G) *C. rodentium* test set and (H) pTox-KatG test set.

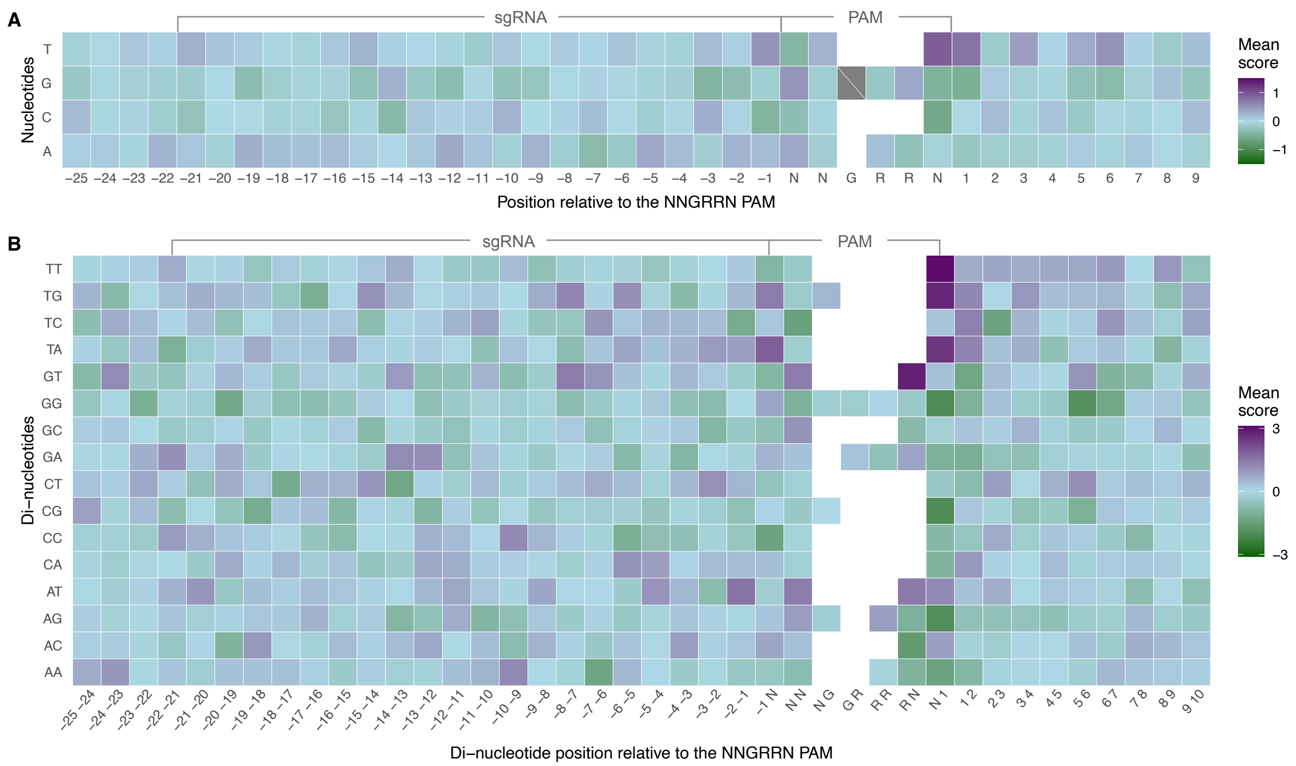


Figure S4: Nucleotide preference across all (Tev)SaCas9/sgRNA target sites in the pTox-KatG enrichment experiment. **(A)** Heatmap of mean single or **(B)** di-nucleotide activity score per position.

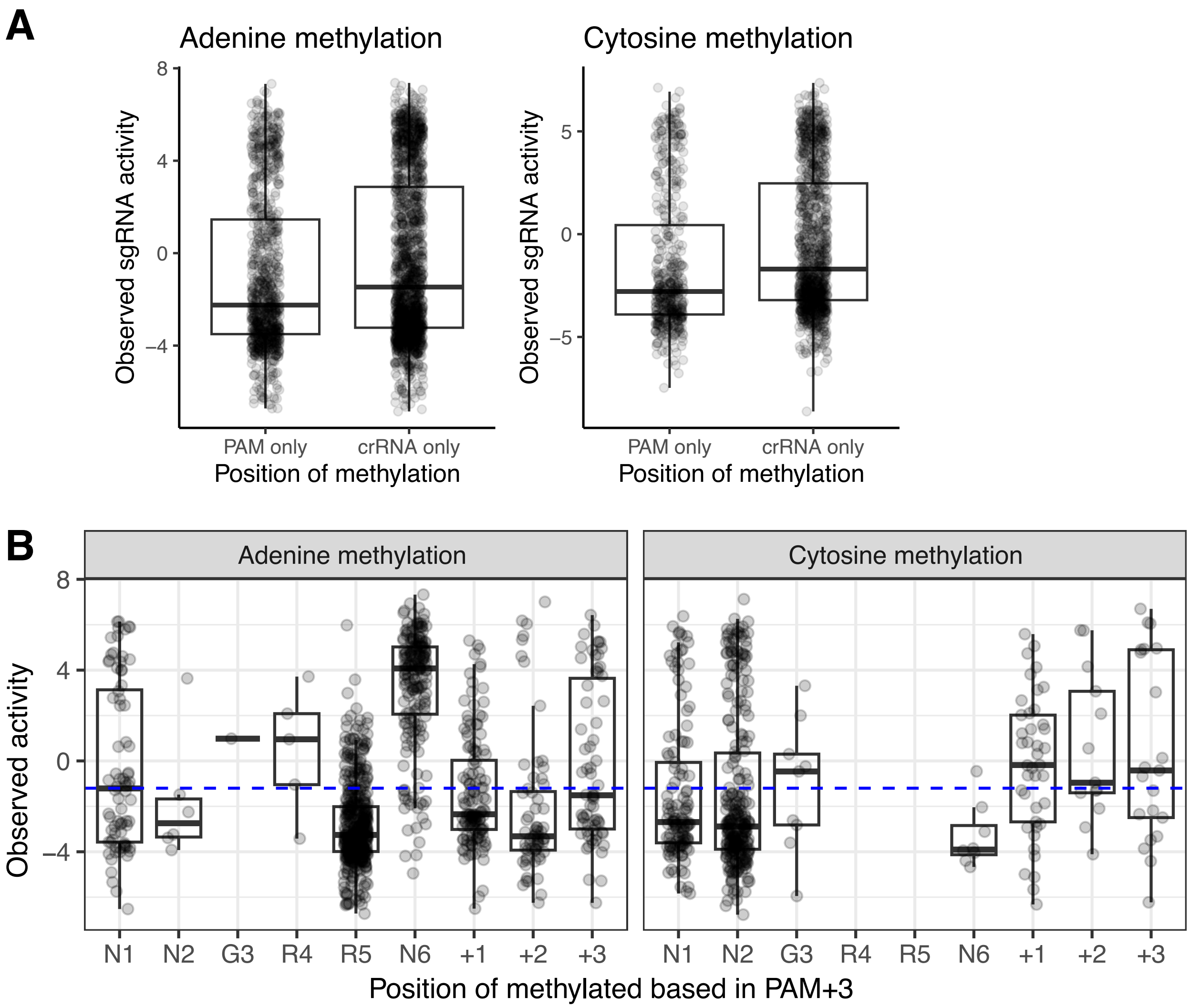


Figure S5: Impact of the position of single adenine and cytosine methylation on sgRNA activity in *C. rodentium*. **(A)** Boxplots of sgRNA activity for single adenine (left) and cytosine (right) methylation events in the PAM region or in the crRNA region. Each point represents the observed activity of a sgRNA targeting a site with the indicated methylation. **(B)** Observed sgRNA activity for adenine or cytosine methylation events mapped to the PAM[+3] region of C. rodentium target sites. Each point represents the activity of an sgRNA with a methylation event at the indicated position. The blue dashed line is the mean observed activity for all sgRNAs tested.

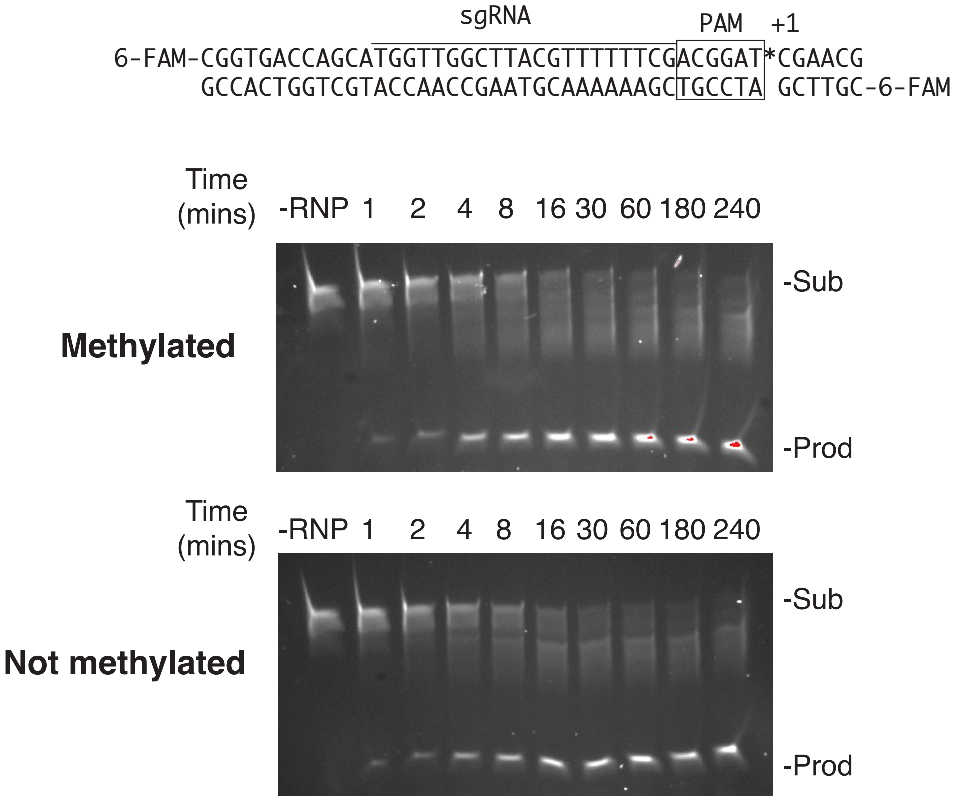


Figure S6: Cytosine (m5C) methylation in the PAM[+1] position does not impact SaCas9 activity on a synthetic DNA substrate. Top, sequence of target site with 6-FAM modifications at the 5′ ends. Bottom, cleavage time course assays with an RNP consisiting of purified SaCas9 and *in vitro* synthesized sgRNAs, or with no added RNP, on the methylated or non-methylated target sites. Aliquots of the stopped reaction were run on a 20% denaturing polyarcylamide gel and visualized on a BioRad Imaging system. The unreacted substrate and products are labeled.

### Supplementary Tables

Table S1: List of oligonucleotides

| Name | Sequence (5′ to 3′) | Notes |
| --- | --- | --- |
| DE5224 | CCCTAAGAAATGAACTGGCAGC | Used in second strand synthesis reaction to make the oligo Pools of sgRNAs double stranded/Reverse primer to amplify *Citrobacter rodentium* sgRNA pool from Twist |
| DE5231 | CCTGGTTCTTGGTCTCTCAC | Forward primer to amplify *Citrobacter rodentium* sgRNA pool from Twist |
| DE7766 | ACACTCTTTCCCTACACGACGCTCTTCCGATCTNNNCAGTCGTTAAGAAGTGATAGAGATACTGAGCACG | Forward primer with Illumina adapter, 3 random nucleotides, 12-mer barcode (CAGTCGTTAAGA), loci-specific nts |
| DE7767 | ACACTCTTTCCCTACACGACGCTCTTCCGATCTNNNCACTACGCTAGAAGTGATAGAGATACTGAGCACG | Forward primer with Illumina adapter, 3 random nucleotides, 12-mer barcode (CACTACGCTAGA), loci-specific nts |
| DE7768 | ACACTCTTTCCCTACACGACGCTCTTCCGATCTNNNGCTCGAAGATTCAGTGATAGAGATACTGAGCACG | Forward primer with Illumina adapter, 3 random nucleotides, 12-mer barcode (GCTCGAAGATTC), loci-specific nts |
| DE7769 | ACACTCTTTCCCTACACGACGCTCTTCCGATCTNNTGAACGTTGGATAGTGATAGAGATACTGAGCACG | Forward primer with Illumina adapter, 2 random nucleotides, 12-mer barcode (TGAACGTTGGAT), loci-specific nts |
| DE7770 | ACACTCTTTCCCTACACGACGCTCTTCCGATCTNNATGGTTCACCCGAGTGATAGAGATACTGAGCACG | Forward primer with Illumina adapter, 2 random nucleotides, 12-mer barcode (ATGGTTCACCCG), loci-specific nts |
| DE7771 | ACACTCTTTCCCTACACGACGCTCTTCCGATCTNNCGAGGGAAAGTCAGTGATAGAGATACTGAGCACG | Forward primer with Illumina adapter, 2 random nucleotides, 12-mer barcode (CGAGGGAAAGTC), loci-specific nts |
| DE7772 | ACACTCTTTCCCTACACGACGCTCTTCCGATCTNTACTACGTGGCCAGTGATAGAGATACTGAGCACG | Forward primer with Illumina adapter, 1 random nucleotide, 12-mer barcode (TACTACGTGGCC), loci-specific nts |
| DE7773 | ACACTCTTTCCCTACACGACGCTCTTCCGATCTNGTTCCTCCATTAAGTGATAGAGATACTGAGCACG | Forward primer with Illumina adapter, 1 random nucleotide, 12-mer barcode (GTTCCTCCATTA), loci-specific nts |

| | | |
|---|---|---|
| DE7774 | ACACTCTTTCCCTACACGACGCTCTTCCGATCTNACGATATGGTCAAGTGATAGAGATACTGAGCACG | Forward primer with Illumina adapter, 1 random nucleotide, 12-mer barcode (ACGATATGGTCA), loci-specific nts |
| DE7775 | CGGTCTCGGCATTCCTGCTGAACCGCTCTTCCGATCTNNNACTCACAGGAATTTTAGTAGATTCTGTTTCCAGAGTAC | Reverse primer with Illumina adapter, 3 random nucleotides, 12-mer barcode (ACTCACAGGAAT), loci-specific nts |
| DE7776 | CGGTCTCGGCATTCCTGCTGAACCGCTCTTCCGATCTNNGTAGGTGCTTACTTTAGTAGATTCTGTTTCCAGAGTAC | Reverse primer with Illumina adapter, 2 random nucleotides, 12-mer barcode (GTAGGTGCTTAC), loci-specific nts |
| DE7777 | CGGTCTCGGCATTCCTGCTGAACCGCTCTTCCGATCTNCAGTCGTTAAGATTTAGTAGATTCTGTTTCCAGAGTAC | Reverse primer with Illumina adapter, 1 random nucleotide, 12-mer barcode (CAGTCGTTAAGA), loci-specific nts |
| DE7820 | GGATCTAGGTGAAGATCCTTTTTGATAATC | Forward primer to amplify pTox+LacIq backbone |
| DE7821 | GGTCTGACGCTCAGTGG | Reverse primer to amplify pTox+LacIq backbone |
| DE7830 | GATTATCAAAAAGGATCTTCACCTAGATCCGGCTGGCCTGGAAGTTGGCATTTCGCTGCT | Forward primer to amplify *Citrobacter* gDNA fragment |
| DE7831 | CATCCTGCATGTTCACCCACTGACGCAGCGTGGTCTGACAGTTACCAATGCTTAATCAGT | Reverse primer to amplify *Citrobacter* gDNA fragment |
| DE6018 | AAAATCTCGCCAACAAGTTGACGAGATAAACACGGCATTTTGCCTTGTTTTAGTAGATTCTGTTTCCAGAGTACTAAAAC | Universal scaffold primer for SaCas9 sgRNAs |
| DE8170 | aagcTAATACGACTCACTATATGGTTGGCTTACGTTTTTTCGGTTTTAGTACTCTGGAAACAG | $sgRNA_{1447}$ for pTox-KatG |
| DE8171 | aagcTAATACGACTCACTATAAGGCCGCGATTAAATTCCAACGTTTTAGTACTCTGGAAACAG | $sgRNA_{295}$ for pTox-KatG |
| DE8802 | aagcTAATACGACTCACTATAGAGCAGATTACGCGCAGAAAAAGTTTTAGTACTCTGGAAACAG | $sgRNA_{4053}$ for pTox-KatG |
| DE8803 | aagcTAATACGACTCACTATAGATCTACGTTAACCCGGAAGGGGTTTTAGTACTCTGGAAACAG | $sgRNA_{1003}$ for pTox-KatG |
| DE8804 | aagcTAATACGACTCACTATAGCAAAGGCTTCATTAAAGGCCTGTTTTAGTACTCTGGAAACAG | $sgRNA_{1543}$ for pTox-KatG |
| DE8805 | aagcTAATACGACTCACTATAGCAGACCCCGTAGAAAAGATCAGTTTTAGTACTCTGGAAACAG | $sgRNA_{4116}$ for pTox-KatG |
| DE8806 | aagcTAATACGACTCACTATAGCAGCAGACGTGCACTGGCCAGGTTTTAGTACTCTGGAAACAG | $sgRNA_{2860}$ for pTox-KatG |

| | | |
|---|---|---|
| DE8807 | aagcTAATACGACTCACTATAGCCGGGCCGGGTGGATGCGCGTGTTTTAGTACTCTGGAAACAG | sgRNA$_{1989}$ for pTox-KatG |
| DE8808 | aagcTAATACGACTCACTATAGCGAGGCTTTATTCGTCGTTTTGTTTTAGTACTCTGGAAACAG | sgRNA$_{1903}$ for pTox-KatG |
| DE8809 | aagcTAATACGACTCACTATAGCTACCAGCGGTGGTTTGTTTGGTTTTAGTACTCTGGAAACAG | sgRNA$_{4031}$ for pTox-KatG |
| DE8810 | aagcTAATACGACTCACTATAGGACGGGAAATCTTCTCGAACTGTTTTAGTACTCTGGAAACAG | sgRNA$_{1505}$ for pTox-KatG |
| DE8811 | aagcTAATACGACTCACTATAGGCACACTGGCGGCCGTTACTAGTTTTAGTACTCTGGAAACAG | sgRNA$_{3227}$ for pTox-KatG |
| DE8259 | 6-FAM/CGTTCGATCCGTCGAAAAAACGTAAGCCAACCATGCTGGTCACCG | Complement of DE8260, synthetic methyl cytosine substrate |
| DE8260 | 6-FAM/CGGTGACCAGCATGGTTGGCTTACGTTTTTTCGACGGAT/iMe-dC/GAACG | Synthetic SaCas9 substrate with 5mC modification in PAM[+1] sequence |

| | | DAM(+) | | DAM(-) | | |
|---|---|---|---|---|---|---|
| **sgRNA:PAM[+1]** | **pTox position** | $k_{obs}$ | **std.dev** | $k_{obs}$ | **std.dev** | **DAM(-) / DAM(+)** |
| 2:ACGGAT[C] | 1447 | 0.19 | 0.03 | 0.63 | 0.22 | 3.32 |
| 2:ACGGGT[C] | 1447 | 0.70 | 0.18 | 0.77 | 0.18 | 1.11 |
| 3:ATGGAT[G] | 295 | 4.63 | 0.78 | 3.76 | 0.94 | 0.81 |
| 3:ATGGAT[C] | 295 | 1.11 | 0.11 | 3.38 | 0.78 | 3.06 |
| 5:AAGGAT[C] | 4053 | 0.41 | 0.19 | 0.51 | 0.33 | 1.24 |
| 6:CCGGAT[C] | 1003 | 0.10 | 0.03 | 1.66 | 0.22 | 16.86 |
| 7:GCGGAT[C] | 1543 | 1.24 | 0.04 | 3.66 | 0.45 | 2.96 |
| 8:AAGGAT[C] | 4116 | 0.68 | 0.20 | 0.86 | 0.41 | 1.26 |
| 9:GGGGAT[C] | 2860 | 0.04 | 0.01 | 0.18 | 0.06 | 4.35 |
| 10:CAGGAT[C] | 1989 | 0.01 | 0.00 | 0.01 | 0.00 | 1.75 |
| 11:CTGGAT[C] | 1903 | 0.08 | 0.03 | 1.31 | 0.12 | 15.72 |
| 12:CCGGAT[C] | 4031 | 0.01 | 0.00 | 0.03 | 0.03 | 3.20 |
| 13:CCGGAT[C] | 1505 | 0.62 | 0.04 | 5.15 | 1.45 | 8.24 |
| 14:AAGGAT[C] | 3227 | 1.33 | 0.12 | 3.69 | 0.64 | 2.77 |

Table S5: Observed reaction rates ($min^{-1}$) for sgRNAs targeting pTox-KatG at the indicated position. The PAM[+1] sequence for each target site is indicated, and substitutions to the sequence are indicated by underlined nucleotides.
